# Psychedelic 5-HT_2A_ receptor agonism: neuronal signatures and altered neurovascular coupling

**DOI:** 10.1101/2023.09.23.559145

**Authors:** Jonah A. Padawer-Curry, Oliver J. Krentzman, Chao-Cheng Kuo, Xiaodan Wang, Annie R. Bice, Ginger E. Nicol, Abraham Z. Snyder, Joshua S. Siegel, Jordan G. McCall, Adam Q. Bauer

## Abstract

Psychedelics hold therapeutic promise for mood disorders due to rapid, sustained results. Human neuroimaging studies have reported dramatic serotonin-2A receptor-(5-HT**_2A_**R)-dependent changes in functional brain reorganization that presumably reflect neuromodulation. However, the potent vasoactive effects of serotonin have been overlooked. We found psilocybin-mediated alterations to fMRI-HRFs in humans, suggesting potentially altered NVC. To assess the neuronal, hemodynamic, and neurovascular coupling (NVC) effects of the psychedelic 5-HT**_2A_**R agonist, 2,5-Dimethoxy-4-iodoamphetamine (DOI), wide-field optical imaging (WFOI) was used in awake Thy1-jRGECO1a mice during stimulus-evoked and resting-state conditions. While DOI partially altered tasked-based NVC, more pronounced NVC alterations occurred under resting-state conditions and were strongest in association regions. Further, calcium and hemodynamic activity reported different accounts of RSFC changes under DOI. Co-administration of DOI and the 5-HT**_2A_**R antagonist, MDL100907, reversed many of these effects. Dissociation between neuronal and hemodynamic signals emphasizes a need to consider neurovascular effects of psychedelics when interpreting blood-oxygenation-dependent neuroimaging measures.

## 1) Introduction

Clinical trials have provided compelling evidence that psychedelics offer rapid and substantial improvement in mood disorders and substance use disorders [1–8]. These remarkable findings have ignited growing interest in elucidating underlying therapeutic mechanisms. Psychedelics, such as psilocybin and lysergic acid diethylamide elicit profound changes in perception, mood, and cognition [9–13], collectively known as the enigmatic “psychedelic experience”. Understanding the neurobiological basis of these effects is a crucial step towards advancing the therapeutic potential of these agents.

Recent high-profile human neuroimaging studies employing functional magnetic resonance imaging (fMRI) have shed light onto the effects of psychedelics on functional brain organization [14–18]. For example, psychedelics have been shown to enhance brain network integration and increased global resting-state functional connectivity (RSFC) [16–20]. These discoveries have led to a burgeoning body of research using fMRI to understand acute and persistent effects of psychedelics at the network level. fMRI indirectly indexes neural activity through changes in blood-flow and metabolism that are reflected in modulations of blood-oxygenation-level-dependent (BOLD) signaling. This linkage is commonly referred to as neurovascular coupling (NVC) [21–26]. Through NVC, changes in RSFC are frequently interpreted as reflecting altered interregional neuronal synchrony. However, it remains unclear whether psychedelic-induced changes in RSFC are a result of neuronal activity, alterations in vascular tone, NVC, or some combination thereof.

The etymology of serotonin (‘serum’ + ‘tone’) reflects its discovery as a substance in blood serum that strongly modulates vascular tone [27]. It was later found to exert potent neuromodulatory effects in the brain, influencing, for example, mood, sleep, and memory [28–31]. Psychedelic-induced hallucinations depend on activity of 5-HT receptors, primarily at the 5-HT**_2A_** receptor (5-HT**_2A_**R) [10, 32–35] and also elicit dose-dependent vasoactive effects [36, 37]. Furthermore, within the central nervous system, 5-HT**_2A_** receptors are expressed not only in neurons but also astrocytes [38] and other glial cells involved in NVC [39–43]. Considered collectively, these observations raise critical questions regarding the proper interpretation of altered neural activity induced by psychedelics as measured by fMRI.

To this end, we first reanalyzed previously published human fMRI data [44] and found modified task-evoked hemodynamic responses in subjects under the acute influence of psilocybin. To assess whether this phenomenon can be attributed to a neuronal, vascular, or neurovascular effect, we employed wide-field optical imaging (WFOI) in awake mice [45]. Our findings suggest that the 5-HT**_2A_**R psychedelic, DOI, substantially altered NVC in a regional– and frequency-specific manner, thereby affecting hemodynamic estimates of underlying neuronal activity (*e.g.*, power spectral density and RSFC). Notably, many of these effects were dose-dependent and reversed by the selective 5-HT**_2A_**R antagonist, MDL. These observations necessitate caution in interpreting the effects of classic psychedelics on neuronal function as assessed through hemodynamic imaging methods, such as fMRI.

## 2) Results

### 2.1) Psilocybin alters stimulus-evoked hemodynamic activity in humans as assessed by FMRI

Previously-published human fMRI data [44] were reanalyzed to evaluate task responses in subjects who received psilocybin, methylphenidate (to control for subjective effects), or no compound while performing a simple auditory-visual matching task (**Fig. 1a**, **Extended Data Fig. 1a**). Generalized linear models were fit for task-evoked activity in left/right visual cortex, left hand, left/right auditory cortex, and left language (Wernicke’s-like area). A double gamma function was employed to model hemodynamic response functions (HRFs), characterized by three-parameters: Peak Value (P), Dispersion (D), and Time to Peak (T). Psilocybin decreased time-to-peak in all regions except the right visual region. Additionally, the dispersion within the left and right visual and auditory regions was decreased, along with peak amplitude in left and right visual regions. These findings suggest a potentially altered relationship between neuronal and hemodynamic activity under acute psychedelic exposure.

**Figure 1).**
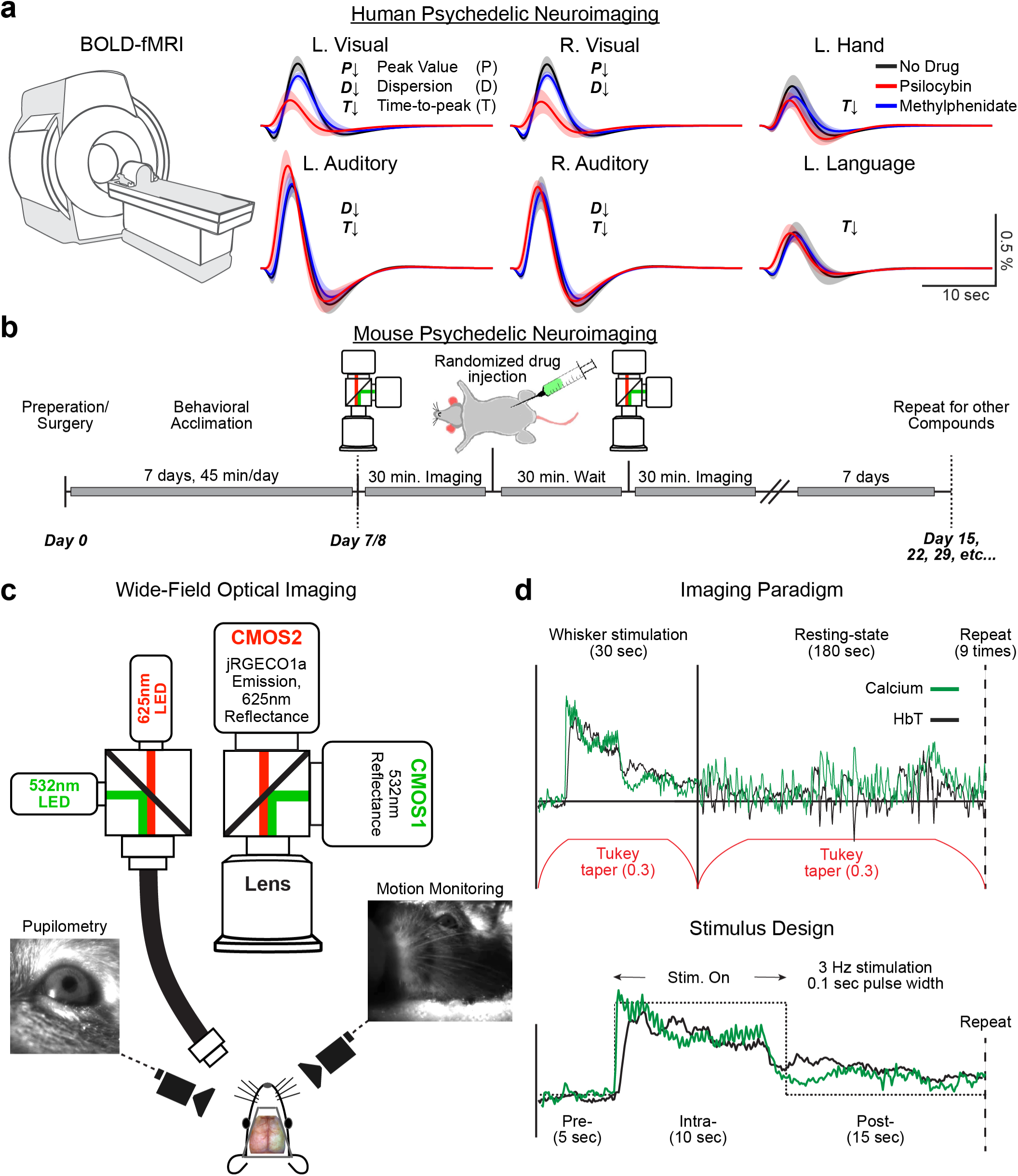
Measuring the effect of 5HT_2A_ receptor agonism on cortical neuronal and hemodynamic activity. a) Psychedelics alter hemodynamic response functions (HRFs) in humans as reported by BOLD-fMRI. Previously published fMRI data from a simple auditory-visual matching task in humans under the acute exposure psilocybin were analyzed [1]. Briefly, subjects were left hand, left/right auditory cortex, and left language (Wernicke’s-like area). HRFs were fit using double gamma basis functions, which were characterized by three-parameters: Peak Value (P), Dispersion (D), and Time to Peak (T). Statistical differences in each parameter across conditions were assessed *via* a 2-way ANOVA; significant effects were evaluated post-hoc *via* t-tests. Data are quantified in Fig. E1a. Psilocybin decreased peak value in left and right visual regions (ANOVA p=0.03 and p=0.020, respectively), decreases time-to-peak in all regions except the right visual region (ANOVA, left visual, p=0.03 left hand, p=0.006; left auditory, p=0.002; right auditory, p=0.044; left language, p=0.007), and decreased dispersion in the left and right visual regions, and left and right auditory regions (ANOVA, p=0.007, p=0.004, p=0.021, p=0.022, respectively). b-e) WFOI in mice. b) Experimental timeline. Prior to imaging, mice were acclimated to awake WFOI for 7 days, 45 minutes per day. Mice were imaged for ∼30 minutes to establish pre-injection metrics, injected with a compound, and after a period of 30 minutes, imaged again for 30 minutes. Compounds were randomized each day of imaging. One week later this process was repeated with a different compound. c) Schematic of WFOI system. WFOI allows for concurrent imaging of calcium and hemodynamic activity over the cortex. A custom light engine consisting of 532 nm and 625 nm LEDs illuminated the skull. Fluorescence emission and diffuse reflectance were collected by a lens, split by a 580 nm dichroic, and sampled by two sCMOS cameras. A 500 nm long pass filter in front of CMOS1 passed 530 nm reflectance while the dichroic and a 590 nm long pass filter in front of CMOS2 passed jRGECO1a emission and 625 nm reflectance. A second 700 nm long pass filter in front of CMOS2 blocked near infrared light used for movement and pupil monitoring. d) Imaging paradigm. Awake whisker stimulation was interwoven with awake resting-state imaging. Ten seconds of air puffs (40 PSI, 100 ms pulses delivered at 3 Hz) were followed by 200 seconds of recovery. The last 180s seconds were considered “resting-state” epochs. This paradigm was repeated 9 times for each imaging session. Stimulus evoked responses and resting-state activity were temporally segmented using a Tukey window (with lobe parameter of 0.3) to minimize boundary effects.

### 2.2 Mapping the effects of hallucinogenic, 5-HT**_2A_**R agonism using wide field optical imaging

Wide-field optical imaging (WFOI) was performed in awake mice expressing the red-shifted genetically encoded calcium indicator, jRGECO1a, under a *Thy1* promotor. This approach enables simultaneously assessment of changes in cortical excitatory neuronal and hemodynamic activity following acute exposure to DOI during both stimulus-evoked and resting-state conditions (**Fig. 1b-d**, **Extended Data Fig. 1b**). The selective 5-HT**_2A_**R antagonist, MDL, was co-administered with DOI to investigate the role of 5-HT**_2A_**R agonism on observed changes. Sub-hallucinogenic doses of DOI and the non-hallucinogenic 5-HT2AR agonist, Lisuride, were used to assess the necessity of hallucinogenic 5-HT**_2A_**R activation for these observations (**Fig. S1**). For all WFOI experiments, raw jRGECO1a fluorescence was corrected for absorption artifacts due to hemoglobin, facilitating concurrent measures of calcium-versus hemodynamic-based neurophysiology (**Fig. S2**). Potential changes in movement during WFOI before and after compound injection were evaluated *via* pupillometry and motion tracking of facial and body movements, which indicated no effect of drug on these measures (**Extended Data Fig. 2** and **Fig. S3**), a finding in line with prior reports [46, 47].

Changes in calcium fluorescence were used as a proxy for neuronal activity, primarily thought to arise from action potential-dependent activation of voltage-gated calcium channels [48]. However, activation of Gq-coupled receptors, such as 5-HT**_2A_**R, can also trigger intracellular calcium release [49]. To assess the potential impact of these signaling cascades on WFOI results, a separate set of slice experiments were performed in Thy1-jRGECO1a mice. Importantly these experiments included positive validation controls of High K+ and NMDA, both of which contribute both substantial action-potential-dependent and action-potential-independent calcium signals. Here, DOI caused a relatively slow and weak action potential-independent increase in calcium fluorescence, suggesting the calcium signal in WFOI during DOI predominantly arises from action potential-dependent mechanisms (**Extended Data Fig. E3**).

### 2.2) 5-HT_2A_ receptor agonism differentially alters stimulus-evoked neuronal and hemodynamic activity and their coupling

#### 2.2.1) Response topographies and dynamics

Before injection of any compound (pre-injection), stimulation of the right whiskers elicited neuronal and hemodynamic activity within left primary somatosensory barrel (S1b) cortex and motor barrel cortex (**Fig. 2a**). DOI differentially modified calcium versus hemodynamic response topographies. For example, while calcium activity exhibited reduced activity in retrosplenial and somatomotor regions, hemodynamic activity was reduced in motor cortex and increased in somatosensory and auditory regions (black arrows in **Fig. 2a**). DOI reduced the maximum neuronal response effect size but did not alter the maximum response amplitude, a finding potentially owing to increased variability in pre-stimulus activity and/or increased intra-stimulus variability (**Extended Data Fig. 1c**). In contrast, hemodynamic response areas and effect sizes were not significantly different within (*i.e.*, pre-versus post-injection) or across compounds (*i.e.*, post-minus post-injection changes across compounds).

**Figure 2).**
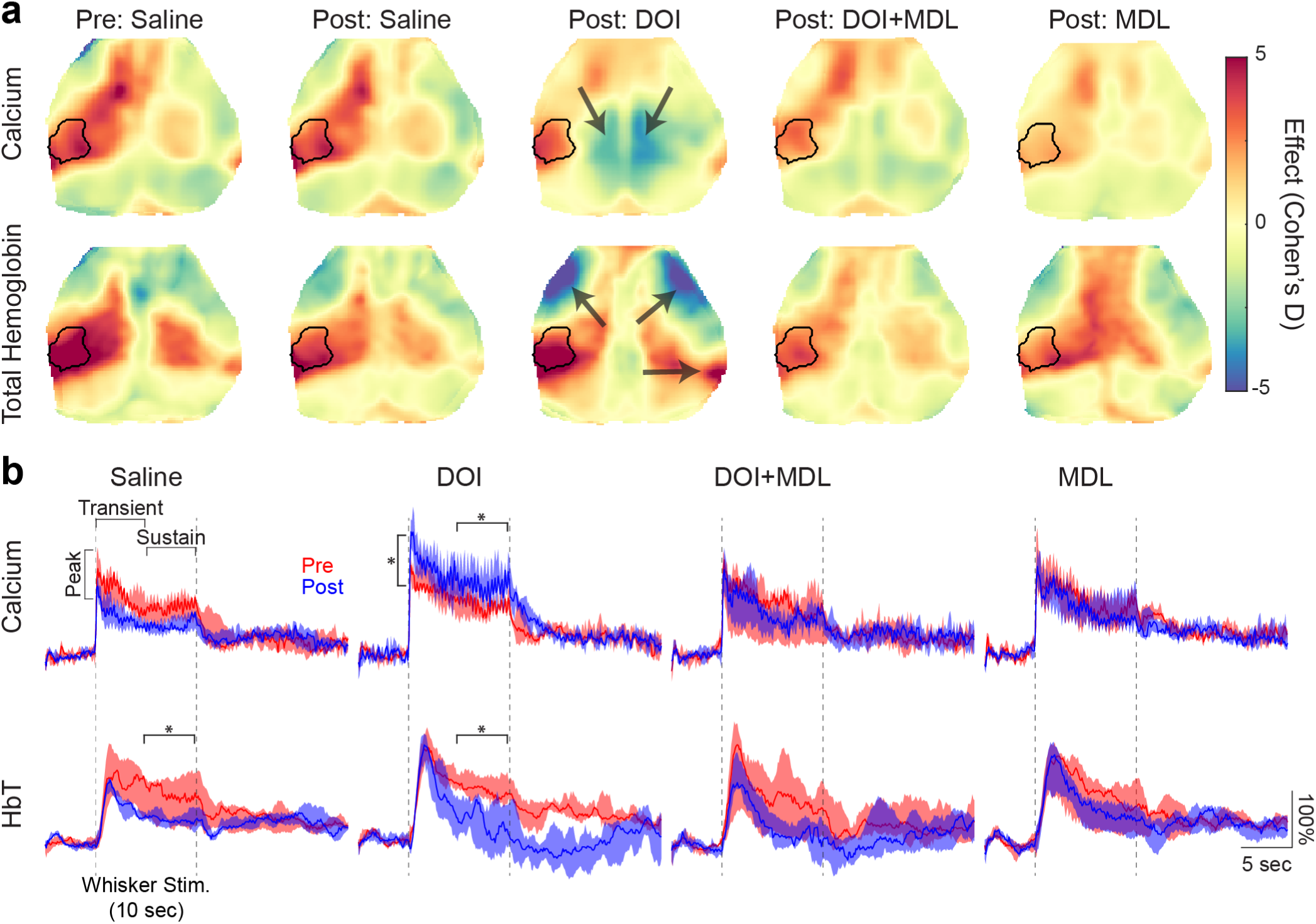
Hallucinogenic 5-HT_2A_ receptor agonism differentially alters stimulus-evoked calcium and hemodynamic activity. a) Response effect maps. Cortical responses to whisker stimulation are reported prior to saline injection and after compound injections. Effect maps were calculated as the Cohen’s D effect size between pre– and intra-stimulus time periods. The black contour represents region of interest (ROI) extracted from pre-injection maps (**Fig. E1b**). DOI decreased calcium response amplitudes within the ROI (–25.72% [–32.4%, –6.52%]; pre-versus post-injection: p=0.032). In contrast, hemodynamic response effects were not differentially altered by compound (Kruskal-Wallis: p=0.643). DOI differentially affected calcium versus hemodynamic response topography (black arrows); there was reduced calcium response in retrosplenial and posterior cingulate regions but decreased hemodynamic response in somatomotor regions and increased hemodynamic response in contralateral auditory cortex. Consequently, DOI decreased the spatial correlation between calcium and hemodynamic response topographies (Δrr = –0.45 [–0.63, –0.28]; Kruskal-Wallis: p=0.009; saline versus DOI: p=0.009). **b) Evoked response dynamics.** Time courses within the ROI were averaged and visualized as medians with shaded error representing the 25^th^ and 75^th^ percentiles. Both calcium and hemodynamic response dynamics were normalized by their average intra-stimulus magnitude before injection to facilitate visual comparison within and across compounds. Therefore, a post injection value of 100% indicates a factor of 2 increase in the intra-stimulus mean compared to the pre-injection mean. Response temporal dynamics were characterized for the first 2 seconds of the whisker response (peak; 0.2-2.2 sec for calcium and 2.0-4.0 sec), 0-5 seconds (transient) and 5-10 seconds (sustained) after stimulus onset. To account for changes in pre– and intra-stimulus variability, Cohen’s D effect size was also computed. DOI increased the peak (26.0% [24.2%, 36.2%]; pre-versus post-injection=0.031) and sustained calcium response (40.0% [3.4%, 120.7%]; pre-versus post-injection: p=0.031), but not the transient response and effect size, suggesting an increase in peri-stimulus variability. In contrast, hemodynamic sustained responses decreased after both DOI (–70.3% [–132.3%, –39.5%]; pre-versus post-injection: p=0.031) and DOI+MDL (–93.6 [–138.5%, =56.6%]; pre-versus post-injection: p=0.031). DOI also decreased the effect size of the evoked hemodynamic response dynamics (Cohen’s D effect of pre-versus post-injection responses in panel b; –38.8% [–63.1%, –24.0%]; pre-versus post-injection: p=0.031). *p<0.05.

Next, to investigate whether DOI alters the temporal dynamics of evoked calcium and hemodynamic activity, time traces within S1b (black contour in **Fig. 2a**) were averaged (**Fig. 2b**). DOI increased both the calcium peak and sustained responses but did not alter the effect size, suggesting increased peri-stimulus variability. In contrast, the sustained hemodynamic response decreased following both DOI and DOI+MDL, with a significant reduction in effect size observed after DOI. Following DOI, hemodynamic responses exhibited a larger (*i.e.*, more negative) post-stimulus undershoot resulting from decreased oxygenation and increased deoxygenation (**Fig. S4a**, **c**; Supplemental Section D1). The post-stimulus undershoot was not present in calcium responses.

#### 2.2.2) Stimulus-evoked neurovascular coupling

The differential effects of DOI on stimulus-evoked neuronal and hemodynamic activity suggests that DOI may alter neurovascular coupling (NVC). NVC was modelled as a causal linear system between evoked calcium and hemodynamic activity and estimated using weighted least squares deconvolution (**Fig. 3a**; model efficacy evaluated in **Fig. S5a, b**; Supplemental Section D2) [50–52]. DOI altered stimulus-evoked NVC, supporting the hypothesis that the observed changes in human HRFs under acute psychedelic exposure might be driven by altered NVC (**Fig. 1a**). These effects were typified by narrower HRFs (decreased FWHM) and increased transduction of frequencies above 0.5 Hz (**Fig. 3**, **Extended Data Fig. 1d**). As observed in the hemodynamic stimulus responses, the HRF under DOI exhibited a larger post-stimulus undershoot compared to pre-injection conditions (**Fig. S4b**, **c**). Notably, many DOI-induced changes in evoked neuronal and hemodynamic activity and NVC were reversed when DOI was co-administered with MDL. MDL alone did not alter evoked activity or NVC (**Figs. 2-3** and **Extended Data Figs. 1c-d**). Together, these findings highlight the crucial role of hallucinogenic, 5-HT**_2A_**R activation on altered stimulus-evoked activity and NVC.

**Figure 3).**
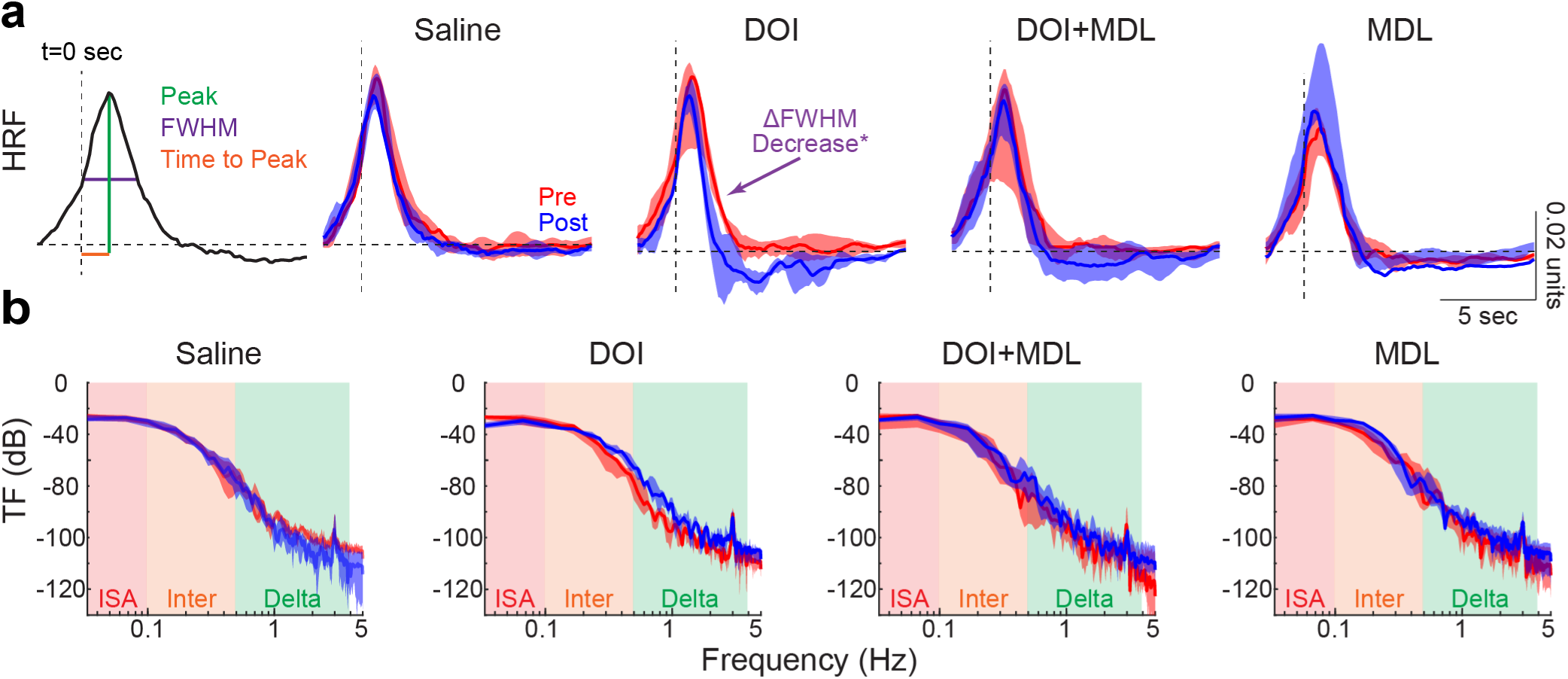
Hallucinogenic 5-HT_2A_ receptor agonism alters stimulus-evoked neurovascular coupling. a) Stimulus-based hemodynamic response functions (HRFs). HRFs were estimated *via* weighted least-squares deconvolution between hemodynamic and calcium response to whisker stimulation. HRFs were parametrized by the peak amplitude (peak, green), time to peak (TTP, orange), and full width at half maximum (FWHM, purple) for each imaging session, mouse, and compound and are presented as medians and 25^th^ and 75^th^ percentiles across mice. DOI significantly decreased the FWHM of the HRF (–31.4% [–36.0%, –27.8%]; pre-versus post-injection: p=0.031; Kruskal-Wallis: p=0.010; saline versus DOI: p=0.028; DOI versus MDL: p=0.006; purple arrow). *p<0.01. The y-axis has units of μMol/(ΔF/F%). **b) Stimulus-evoked transfer functions.** Transfer functions were calculated as the power spectral density estimate of the HRF and are presented as medians and 25^th^ and 75^th^ percentiles across mice. A trend of increased transduction in frequencies above 0.5 Hz was observed for DOI conditions only (delta band increase +367% [57%, 472%]; pre-versus post-injection: p=0.063). Data are presented as medians and 25^th^ and 75^th^ percentiles across mice.

### 2.3) DOI differentially alters resting-state neuronal and hemodynamic activity

Functional neuroimaging under the task-free state has provided key insights into uncovering large-scale patterns of spontaneous brain dynamics and organization that support function [53, 54]. Psychedelics have been shown to markedly alter both regional neuronal synchrony [55–59] and brain-wide hemodynamic activity [16–20]. However, the relation between large-scale changes in spontaneous neuronal and hemodynamic activity induced by psychedelics remains unclear.

Power spectral density estimates (PSDE) of pre-injection global neuronal and hemodynamic activity exhibited a 1/frequency-like profile (**Fig. 4a**,**b**) in accordance with previous mouse studies [45, 60–62]. Following injection of saline, broad-band reductions in neuronal activity were observed (0.16-5.12 Hz; **Extended Data Fig. 4a**). While DOI decreased some low-frequency calcium activity (0.16-0.64 Hz), it led to the emergence of a prominent increase in delta band activity (0.32-5.12 Hz). Notably, hemodynamics under DOI also exhibited increased power in the delta band; however, this effect was narrower in frequency and shifted to lower frequencies (0.32-0.64 Hz; **Extended Data Fig. 4b**). Many of these changes were mitigated when DOI was co-administered with MDL (**Fig. 4** and **Extended Data Fig. 4**). Similarly, MDL alone did not appreciably alter cortical neuronal and hemodynamic power compared to saline.

**Figure 4).**
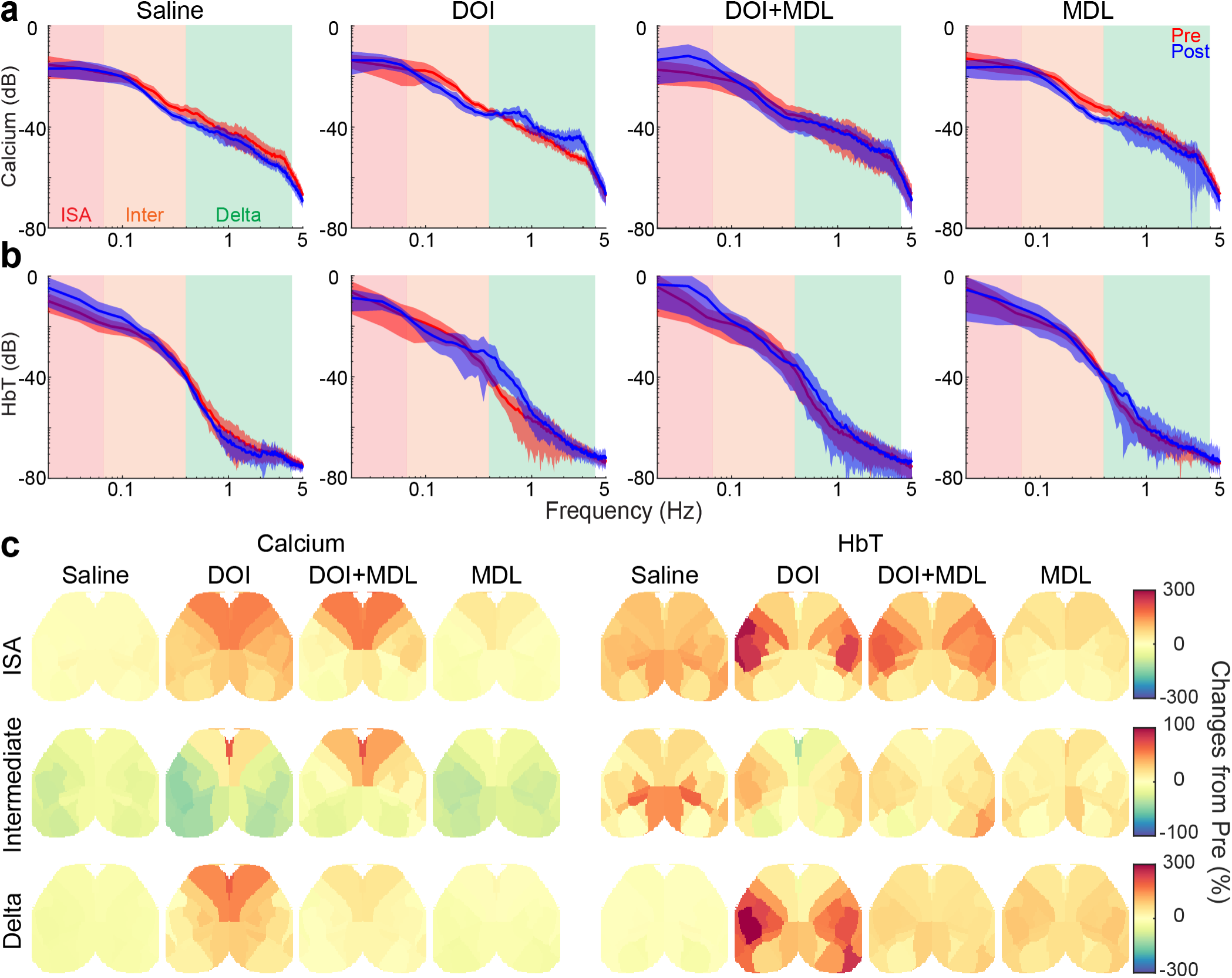
Hallucinogenic 5-HT_2A_ receptor agonism differentially alters resting-state neuronal and hemodynamic activity in a region-dependent manner. Power spectral density estimates (PSDE) were computed for each cortical region using a smoothed estimator (pwelch). Global PSDEs were computed as the mean PSDE across all regions. **a) Global PSDEs of calcium activity.** Data are presented as means and standard deviations across mice. Plots are color-coded according to the following frequency ranges: infraslow activity (ISA), 0.01 to 0.08 Hz, intermediate (inter): 0.08 to 0.5 Hz, and delta 0.5 to 4.0 Hz (global PSDE was also integrated over double octave, partially overlapping frequency bins, see **Fig. E4a-b**). After DOI injection, calcium power decreased between 0.08 Hz and 0.64 Hz (0.08 Hz-0.32 Hz: pre-versus post-injection p=0.009; 0.16 Hz-0.64 Hz: pre-versus post-injection: p=0.020), but increased above 0.32 Hz (ANOVA, 0.32 Hz-1.28 Hz: p=0.0006; 0.64 Hz-2.56 Hz: p=0.0018; 1.28 Hz-5.15 Hz: p=0.0017; saline versus DOI post-hoc p=0.0005, p=0.0007, p=0.0013). DOI+MDL largely preserved pre-injection power spectral characteristics (increases observed only between 0.02 Hz-0.08 Hz: versus post-injection: p=0.018; 0.04 Hz-0.16 Hz: versus post-injection: p=0.008). **b) Global PSDEs of hemodynamic activity.** Following saline injection hemodynamic power increased in lower frequencies (0.02 Hz-0.08 Hz: versus post-injection: p=0.045; 0.04 Hz-0.16 Hz: versus post-injection: p=0.044). DOI increased lower delta band frequencies (0.32 Hz-1.28 Hz: versus post-injection: p=0.002; ANOVA: p<0.0001; post-hoc DOI vs saline: p<0.0001; 0.64 Hz-2.56 Hz: versus post-injection: p=0.006; ANOVA: p<0.0001; post-hoc DOI vs saline: p<0.0001). **c) Regional changes in cortical power.** Band-limited power is displayed as fractional changes from baseline (*i.e.*, pre injection). DOI increased calcium power (*Left*) in the anterior-to-posterior and lateral-to-medial 832 directions: DOI increased ISA power in cingulate, frontal, and secondary motor networks (164% [156%, 186%]) but only slightly increased ISA power in somatosensory and visual networks (26% [17%, 57%] and 13% [10%, 17%], respectively). This feature was attenuated under DOI+MDL, for instance, in cingulate and frontal networks (124% [94%, 157%]). Intermediate activity exhibited similar spatial changes as ISA but significantly smaller in magnitude. For instance, DOI and DOI+MDL minimally increased cingulate (68% and 69%), frontal (33% and 42%), and secondary motor networks (14% and 38%). DOI increased delta-band power in medial regions such as cingulate (197%) and secondary motor regions (153%). DOI+MDL largely reversed these changes (maximum of 49%). Hemodynamic activity (*<u>Right</u>*) reveal differential alterations compared to calcium. Under DOI, ISA appreciably increased over somatosensory cortex (212% [127%, 237%]), an effect attenuated in the DOI+MDL condition (162% [154%, 178%]). Under both DOI and DOI+MDL, changes in intermediate activity reflected those over ISA but approximately three times smaller. Notably, increased delta-band activity was observed under DOI in hemodynamic activity over somatosensory and portions of motor and visual regions, with a 314% increase in primary somatosensory barrel cortex. Full-band, non-integrated calcium and hemodynamic PSDE for each region and compound are displayed in **Fig. E5.**

Given the observed changes in global power, we next examined whether these effects were region-specific (**Figs. 4c**, **Extended Data Fig. 5**). PSDEs for each cortical region defined by the Allen mouse brain atlas were integrated over three frequency ranges (Infraslow activity, ISA: 0.01-0.08 Hz, Intermediate: 0.08 – 0.50 Hz, and delta: 0.5-4.0 Hz, Fig. 4c) and plotted as a percent change from pre-injection. Saline and MDL had minimal effects on calcium band-limited power and slightly larger effects on hemodynamics. The emergent delta-band calcium changes under DOI were largest in cingulate, secondary motor, and other more medial regions (**Fig. 4c**, bottom row). In contrast, the delta-band hemodynamic activity was primarily observed in somatosensory cortex.

Discordant regional changes in neuronal and hemodynamic activity were not limited to the delta band; alterations in infraslow and intermediate calcium activity often occurred in different cortical areas than hemodynamic activity (**Figs. 4, Extended Data Fig. 5**). For instance, DOI-induced changes in intermediate calcium power frequently opposed hemodynamic changes in several regions; calcium activity in the visual, parietal, and somatosensory regions decreased, while hemodynamic activity in those areas remained unchanged or increased. As observed in stimulus conditions, co-administration of DOI and MDL largely reversed the regional effects observed with DOI alone (**Fig. 4c**).

### 2.4) DOI alters resting-state neurovascular coupling

Following our observations of compound-and-region-specific changes in neuronal and hemodynamic activity, we investigated whether 5-HT**_2A_**R agonism differentially affects global NVC (**Figs. 5a-b** and **Extended Data Fig. 4c**) or specifically alters regional NVC (**Figs. 5c** and **Extended Data Fig. 6**).

**Figure 5).**
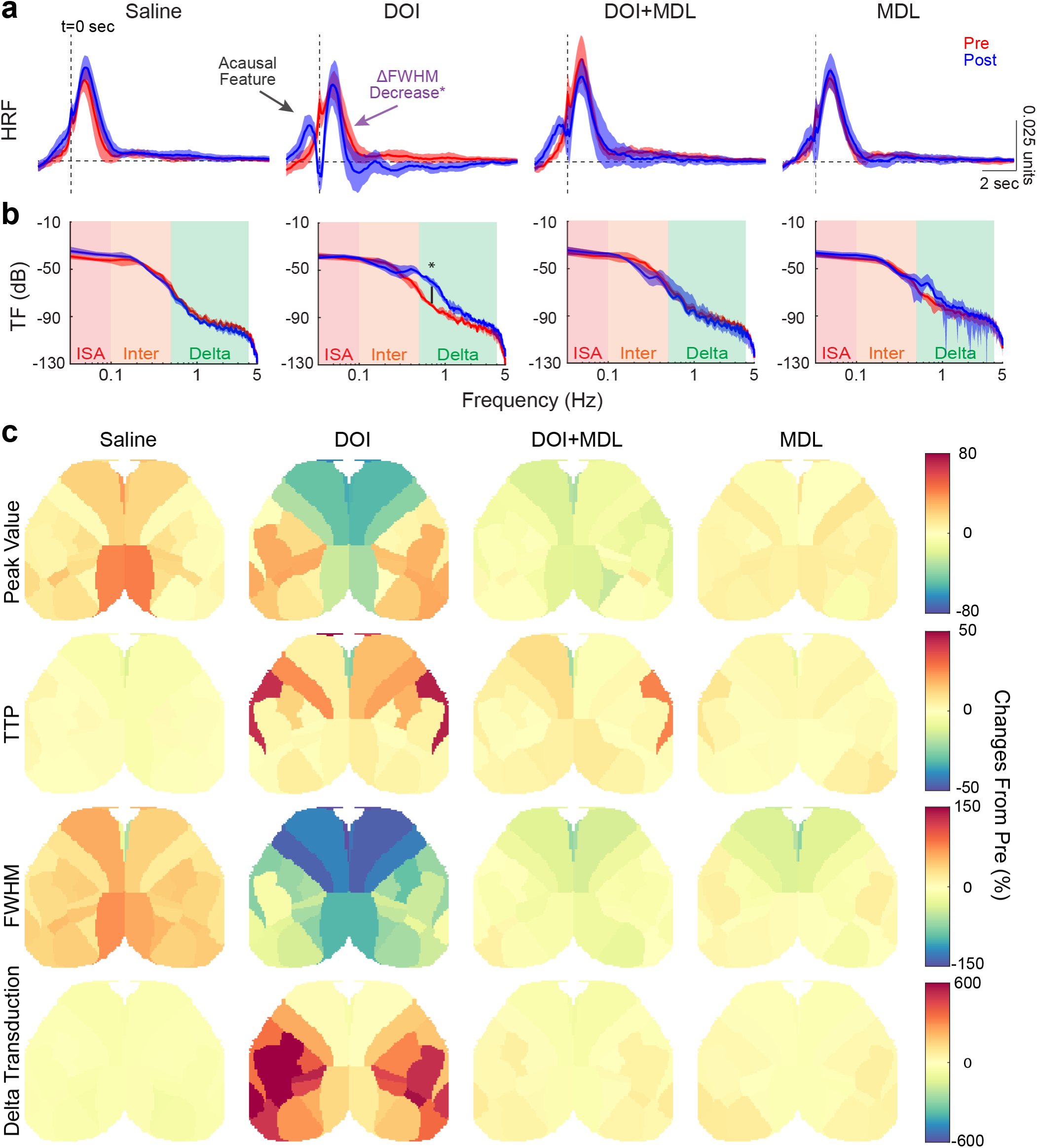
Hallucinogenic 5HT_2A_ receptor agonism alters neurovascular coupling at local and global scales. a) Resting-state hemodynamic response functions. Pre-injection resting-state hemodynamic response functions (HRFs) resembled those observed during whisker stimulation. After DOI injection, HRFs deviated from the canonical form, exhibiting a peak before time zero (black arrow), suggesting an acausal or retrograde (vascular-to-neural relation) relation. HRFs were parameterized by the peak value, time to peak (TTP), and full-width-at-half maximum (FWHM). DOI increased the TTP (ANOVA: p=0.020; saline versus DOI: p=0.019), reduced model fit (saline: from 0.80±0.04 to 0.76±0.05; DOI from 0.76±0.05 to 0.70±0.05; ANOVA: p=0.040; saline versus DOI: p=0.048), and decreased the FWHM (purple arrow, –42.7%±71.1%, pre-versus post-injection: p=0.015; ANOVA: p=0.020; saline versus DOI: p=0.018). The y-axis has units of μMol/ (ΔF/F%). Data are presented as 896 means +/-standard deviations across mice. **b) Resting-state transfer functions.** Transfer functions were calculated as the power spectral density estimates of the HRF and are presented as mean ± standard deviation. DOI induced a quasi-oscillatory feature in lower delta frequencies (321.9%±275.2%: p=0.018; ANOVA: p=0.004; saline versus DOI: p=0.004; DOI versus DOI+MDL: p=0.028). TFs have units of (μMol/(ΔF/F%))^2^/Hz). Data are presented as means +/-standard deviations across mice. **c) Regional neurovascular coupling.** DOI induced focal changes in NVC (Peak, TTP, FHWM, Delta Transduction) of opposite polarity in higher order brain regions (*e.g.*, frontal, and cingulate) versus lateral somatosensory and parietal regions. Coupling amplitude decreased (lower peak values) in frontal/cingulate regions (–62%), while more modest increases occurred in somatosensory regions (*e.g.*, barrel cortex: 32%). DOI delayed NVC (increased TTP) in frontal (51%) and anterior-lateral parts of somatomotor regions (32%). Most strikingly, DOI substantially decreased the FWHM of the HRF over most of the cortex with the largest changes occurring in cingulate (–142%) and frontal (–183%) regions. DOI substantially increased delta-band transduction over most of the cortex (cortical-wide: 237% [79%, 350%], 793% in primary somatosensory barrel cortex), except in cingulate regions where it was reduced by 56%.

Prior to administration of any compound, global HRFs under resting-state conditions closely resembled stimulus-evoked HRFs (**Fig. 5a**) and reflect both our previous findings [63] and those of others [51, 52]. However, after DOI administration, the HRF changed significantly, exhibiting a prominent peak prior to time zero, suggesting a deviation from the assumed causal neuronal-to-hemodynamic relation. This emergent peak gave rise to a quasi-oscillatory feature, manifesting as increased delta-band transduction in the transfer function (TF; **Fig. 5b**). These characteristics were also observed in the lagged cross-covariance analysis (**Extended Data Figs. 7 and 8**, Supplemental Section N1). Moreover, HRF features after time zero (neuronal-to-hemodynamic causality) were also significantly affected by DOI, including increased time-to-peak (TTP) and decreased full-width-at-half-maximum (FWHM; **Extended Data Fig. 4c**). Importantly, DOI also reduced model fit (**Fig. S5b**, **c**), indicating a deviation from the model’s assumptions of causality and linearity.

Recent studies have highlighted that NVC exhibits regional variation across the cortex, even in healthy subjects [64–68]. To determine where on the cortex NVC changes, regional HRFs and TFs were computed (**Extended Data Fig. 6**), parameterized, and visualized (**Fig. 5**). Consistent with these findings, pre-injection HRFs varied across the cortex under saline (**Figs. 4c** and **Extended Data Fig. 6a**, left).

Critically, after injection of DOI, HRFs deviated from canonical form (*i.e.*, presence of a dominant quasi-acausal peak) ubiquitously across the cortex (**Extended Data** Fig 6a, middle). As a result, HRF width deceased across most the cortex, with the most substantial decreases in the cingulate, frontal, and motor regions (**Fig. 5c**, FWHM, **Extended Data Figs. 6a and 8**). Delta band transduction increased within most of visual, parietal and somatosensory cortex and modestly decreased in cingulate (**Fig. 5c, Fig. Extended Data Figs. 6b and 8**). Additionally, DOI often induced alterations in NVC (Peak, TTP, FWHM) that exhibited opposite polarity between higher-order regions (*e.g.*, frontal and cingulate) and lateral somatosensory and parietal regions (**Fig. 5c**). For example, under DOI, the HRF peak decreased within primary and secondary motor and some higher-order brain regions (frontal, cingulate and retrosplenial) but increased within lateral somatosensory and parietal regions (**Fig. 5c**, Peak value); DOI increased HRF delays (larger TTP) within secondary somatosensory and primary motor, but shortened delays in cingulate cortex (**Fig. 5c**, TTP).

We next investigated the degree to which the observed changes in resting-state NVC depended on hallucinogenic versus non-hallucinogenic 5-HT**_2A_**R agonism (**Extended Data Fig. 9**). Sub-hallucinogenic doses of DOI (**Extended Data Fig. 9 a-c**) did not significantly impact global (**Extended Data Fig. 9b**) or local (**Extended Data Fig. 9c**) NVC compared to saline. Separately, we tested the non-hallucinogenic, 5-HT2**_A_**R agonist, Lisuride, selected for its ability to target similar signaling pathways to classic serotonergic psychedelics that may influence vascular tone, without inducing hallucinations at a dose known to alter locomotor behaviors [69] (**Extended Data Fig. 9a**). Lisuride affected cortical calcium and hemodynamic activity similarly to saline, for example, broad-band reductions in neuronal activity across infraslow and intermediate frequency ranges, with more modest effects on hemodynamic signaling (**Extended Data Fig. 9d-e**). Unlike hallucinogenic doses of DOI, the shape of the HRF and the topographies of NVC pre– and post-Lisuride remained preserved and were not different than changes induced by saline (**Extended Data Fig. 9f**, g).

### 2.5) DOI differentially alters functional brain organization

Dramatic alterations in functional organization are a primary focus in psychedelic studies employing large-scale neuroimaging techniques, such as fMRI [14–18]. These network-level resting-state functional connectivity (RSFC) alterations, particularly in regions associated with higher-order cognitive processes [16, 20, 34, 70–72], are often attributed to neuronal origins (*e.g.*, ‘neural correlates’ [20, 34, 73, 74]), with assertions that these alterations contribute to the subjective effects of psychedelics [34, 73–75]. However, considering our observations of significant alterations in neurovascular coupling, it remains unclear whether these changes indicate a reorganization of neuronal network connectivity, reflect modifications in neurovascular coupling, or represent a combination of both.

To align with human neuroimaging analyses, infraslow activity (ISA) RSFC in mice was evaluated for frontal, cingulate, retrosplenial (“higher order” brain regions in mice), and secondary motor (as a control) regions (**Fig. 6**). Prior to injection (**Fig. 6a-d**, top), both calcium– and hemodynamic-based RSFC maps exhibited topographies consistent with prior observations by us and others [60, 61, 76–79], including strong intra-network connections (indicated by high correlations both within and across hemispheres) and network-specific anticorrelations, primarily along the anterior-posterior axis for the frontal, retrosplenial, and secondary motor regions.

**Figure 6).**
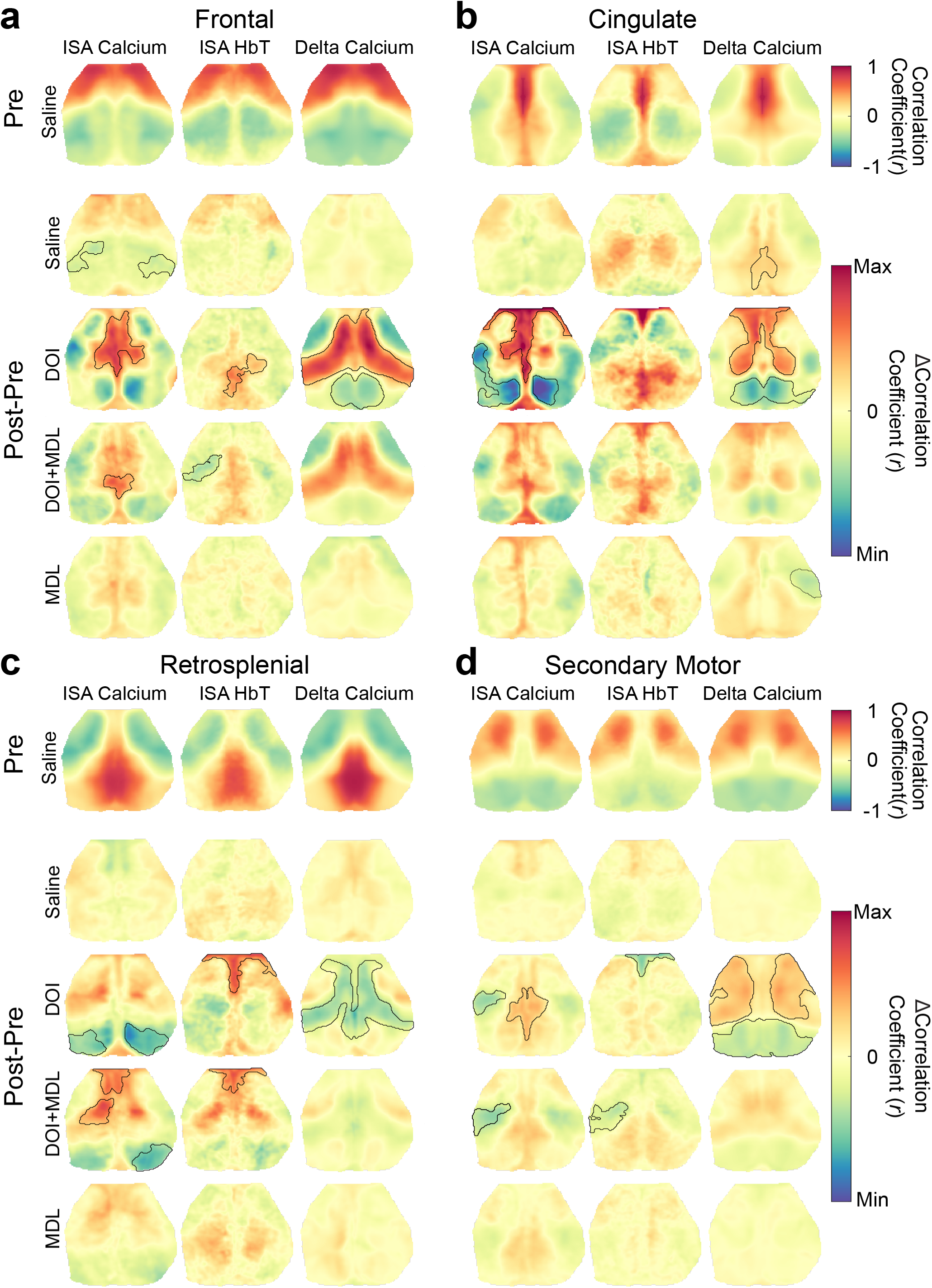
Calcium and hemodynamic activity report different accounts of RSFC changes from hallucinogenic 5-HT2AR agonism. Seed-based resting-state functional connectivity (RSFC) was computed using zero-lag correlations over infraslow (ISA, 0.01 Hz – 0.08 Hz) and delta band (0.5 Hz – 4.0 Hz) activity. To demonstrate the effects of altered NVC on measures of functional organization, RSFC was evaluated in regions exhibiting the largest NVC changes (Frontal, Cingulate and Retrosplenial) and secondary Motor (M2) as a control. **a) Frontal RSFC:** Saline modestly decreased *<u>ISA calcium</u>*functional connectivity between frontal and somatosensory (left, Δrr = –0.13±0.02; right. Δrr = –0.12±0.02). DOI enhanced inter-network *<u>ISA calcium</u>* RSFC between frontal and cingulate (Δrr = +0.31±0.15) and *<u>ISA HbT</u>* connectivity between frontal and posterior retrosplenial (Δrr = +0.18±0.10). DOI increased *<u>delta calcium</u>* RSFC between frontal and secondary motor/posterior somatosensory (Δrr = +0.18±0.03) and increased anticorrelations between frontal and retrosplenial (Δrr = –0.11±0.05). MDL attenuated DOI effects, excluding *<u>ISA calcium</u>* RSFC between frontal and retrosplenial (Δrr = +0.20±0.13) and *<u>ISA HbT</u>* RSFC between frontal and somatosensory (Δrr = –0.16±0.06). **b) Cingulate RSFC:** DOI significantly enhanced *<u>ISA calcium</u>* anticorrelations between cingulate and posterior-lateral retrosplenial (Δrr = –0.41±0.17) and strengthened *<u>ISA calcium</u>* connectivity between cingulate/frontal and anterior-medial retrosplenial regions (Δrr = +0.40±0.24). These DOI-induced *<u>ISA calcium</u>* RSFC changes were not reflected in *<u>ISA HbT</u>* RSFC. DOI increased anticorrelated *<u>delta calcium</u>* activity between cingulate and retrosplenial (Δrr = –0.14±0.07) and increased *<u>delta calcium</u>* connectivity between cingulate and frontal/medial somatosensory (Δrr = 0.15±0.04). MDL reversed all DOI effects. **c) Retrosplenial RSFC:** DOI decreased *<u>ISA calcium</u>* RSFC within retrosplenial and between retrosplenial and lateral visual (left hemisphere: Δrr = –0.22±0.14, right hemisphere: Δrr = –0.31±0.19). DOI also enhanced *<u>ISA HbT</u>* RSFC between retrosplenial and frontal/cingulate (Δrr = +0.29±0.12). MDL did not fully reverse the effects of DOI (*<u>ISA calcium</u>* RSFC between retrosplenial and frontal/cingulate Δrr = 0.25±0.20; between retrosplenial and medial somatosensory Δrr = 0.25±0.16, and between retrosplenial and lateral parts of visual cortex Δrr = –0.28±0.20. *<u>ISA HbT</u>* RSFC between retrosplenial and frontal/cingulate Δrr = 0.26±0.16.). DOI increased *<u>delta calcium</u>* anticorrelations between retrosplenial and M2/posterior somatosensory (Δrr = –0.11±0.02). **c) Secondary Motor RSFC:** DOI increased negative *<u>ISA calcium</u>* RSFC between M2 and lateral somatosensory (Δrr = – 0.18±0.12), and increased connectivity between M2 and anterior cingulate/posterior retrosplenial (Δrr = –0.15±0.07). This effect was not observed in *<u>ISA HbT</u>* RSFC (decreased connectivity between M2 and cingulate of Δrr = –0.20±0.11). *<u>Delta calcium</u>*RSFC under DOI was increased within sensorimotor regions (left hemisphere: Δrr = +0.09±0.03, right hemisphere: Δrr = +0.09±0.05) and increased anticorrelated activity in M2 and retrosplenial/visual (Δrr = +0.09±0.04). MDL administered with DOI reversed many DOI effects, excluding decreases in *<u>ISA calcium</u>* and *<u>ISA HbT</u>* connectivity between M2 and somatosensory cortex. MDL alone slightly decreased connectivity between cingulate and right somatosensory regions (Δrr = –0.09±0.13). The minimum and maximum values for ISA FC maps and delta FC maps are |*r*|<0.5 and |*r*|<0.3, respectively. Inter and Intra RSFC is further evaluated in **Fig. E10**. Statistical differences (black contours) in seed-based RSFC were evaluated on a cluster-wise basis.

After the injection of DOI, cortical-wide neuronal RSFC was profoundly altered. DOI enhanced intra-network calcium RSFC within the cingulate region (**Fig. 6b**) and inter-network calcium RSFC between cingulate and frontal regions (**Fig. 6a-b**). Additionally, DOI significantly altered calcium retrosplenial RSFC across the cortex as follows: (1) enhanced negative connectivity between the cingulate and posterior-lateral parts of retrosplenial regions (**Fig. 6b**) and (2) strengthened connectivity between cingulate/frontal and anterior-medial parts of retrosplenial regions (**Fig. 6a-b**). Furthermore, DOI significantly reduced connectivity between the retrosplenial cortex and portions of the visual network (**Fig. 6c**). These prominent differential alterations in inter-versus intra-network connectivity were further investigated through community detection analysis (**Extended Data Fig. 10**; Supplemental Sections N2). DOI markedly decreased calcium silhouette scores (indicating weakened intra-network connections and strengthened inter-network connections) across most of the cortex, with the most significant decrease observed in the retrosplenial region. These alterations were substantially mitigated in the hemodynamic scores. While DOI+MDL markedly attenuated most of the calcium RSFC changes induced by DOI alone, retrosplenial connectivity remained disrupted in the frontal, anterior sensorimotor, and visual regions (**Fig. 6c**).

Strikingly, almost none of the RSFC changes evaluated by neuronal calcium activity were reflected in hemodynamic measures. For instance, DOI induced only small increases in inter-network hemodynamic RSFC between the frontal and retrosplenial regions, as well as in connectivity between the retrosplenial and anterior cingulate regions. To quantify these topographical disparities between calcium– and hemodynamic-based RSFC, we employed spatial correlation and regression analyses of the RSFC maps (**Fig. 6**; **Table 1**). Briefly, DOI consistently decreased both the correlation and proportionality (*i.e.*, β, as defined in Eq. 5) of the linear relationship between the two measures. As observed in calcium-based RSFC, MDL attenuated most of the hemodynamic RSFC changes induced by DOI, restoring correspondence between neuronal-hemodynamic RSFC (**Table 1**).

**Table 1).** Spatial similarity between RSFC changes estimated by calcium and hemodynamic signaling. Differences (*i.e.*, post-minus pre-injection) in ISA (*i.e.*, 0.01 Hz–0.08 Hz) RSFC topography were computed for each mouse, compound, and region (**Fig. 6**). Spatial similarity (Pearson rr spatial correlation) and linear regression (slope and intercept) evaluated the linear, topographical correspondence between calcium– and hemodynamic-based RSFC. DOI significantly dissociated (*i.e.*, decreased correlation) neuronal versus hemodynamic reports of RSFC in retrosplenial (Kruskal-Wallis, p=0.028) and secondary motor (Kruskal-Wallis, p=0.038) regions. There was also a trend of dissociation in the frontal region. Interestingly, MDL alone decreased RSFC correspondence more than DOI+MDL in the cingulate and retrosplenial region, but increased correspondence more than MDL in frontal and secondary motor regions. Significance was determined using Kruskal-Wallis tests and evaluated post-hoc using paired Wilcoxon sign-rank test and corrected for multiple comparisons.

|  | Frontal | Cingulate | Retrosplenial | M2 |
| --- | --- | --- | --- | --- |
| Saline | y=0.72x+0.01;<br>r=0.61 | y=0.06x+0.03;<br>r=0.05 | y=0.63x+0.01;<br>r=0.49 | y=0.78x-0.02;<br>r=0.58 |
| DOI | y=0.18x+0.00;<br>r=0.29 | y=0.04x+0.05;<br>r=0.05 | y=0.04x+0.02;<br>r=0.05 | y=0.21x-0.02;<br>r=0.25 |
| DOI+MDL | y=0.24x+0.00;<br>r=0.37 | y=0.37x+0.02;<br>r=0.45 | y=0.54x+0.02;<br>r=0.56 | y=0.50x+0.00;<br>r=0.41 |
| MDL | y=0.34x-0.01;<br>r=0.43 | y=0.17x+0.01;<br>r=0.18 | Y=0.42x+0.02;<br>r=0.42 | y=0.61x-0.01;<br>r=0.59 |

Considered collectively, these results suggest that baseline RSFC evaluated through calcium and hemodynamic signals is, in many cases, topographically similar, but quantitatively distinct. Administration of DOI results in substantial disparities between hemodynamic-based measures of RSFC versus underlying neuronal activity which were nearly reversed when DOI was co-administered with MDL.

#### 2.5.1) Calcium measures of RSFC: Delta band

Another advantage of calcium-based measures of RSFC is their ability to capture faster neuronal dynamics, which exhibit diverse features compared to infraslow hemodynamics [63]. Prior to injection, delta band (0.5 Hz-4.0 Hz) calcium activity reveals distinct functional topography compared to ISA RSFC, in line with prior work [60, 77, 79] (**Fig. 6a-d**, right). As observed with ISA RSFC, DOI modified delta band RSFC, most notably affecting intra– and inter-network connectivity (**Extended Data Fig. 10**), with a widespread decrease in retrosplenial connectivity with the rest of the cortex (**Fig. 6a-d**). Moreover, cingulate connectivity increased with frontal and medial motor/anterior lateral somatosensory regions and parietal regions (**Fig. 6a**).

Interestingly, the observed changes in delta RSFC for frontal and retrosplenial regions were topographically similar, but of opposite sign, excluding decreased RSFC with retrosplenial regions (**Fig. 6c** compared to **Fig. 6b**).

Retrosplenial and cingulate cortex comprise aspects of the default mode network in mice and are homologous structures to the posterior and anterior cingulate cortex in humans. The severing of connectivity between these regions is especially relevant given common observations of psychedelics on inter– and intra-network connectivity in humans [16, 18, 80–82]. To that end, we evaluated inter-versus intra-network RSFC (**Fig. E10**; Supplemental Section N2). Briefly, under DOI, the changes in scores were smaller than those observed in ISA, with the largest increases occurring in the retrosplenial cortex. Notably, the most prominent changes in silhouette scores were found at the boundaries of functional parcels, indicating alterations to parcel boundaries.

## 3.) Discussion

### 3.1) Summary of present findings

We demonstrate that previously published human fMRI data reveals altered stimulus-evoked hemodynamic responses in subjects acutely experiencing psilocybin. To assess whether this phenomenon can be attributed to a neuronal, vascular, or neurovascular effect, we employed wide-field optical imaging (WFOI) in awake mice expressing the genetically encoded calcium indicator, jRGECO1a, in cortical pyramidal cells to assess the effects of the psychedelic, DOI, on stimulus-evoked and resting-state neuronal and hemodynamic activity. DOI differentially altered neuronal versus hemodynamic activity both during whisker stimulation and in the resting-state. These observations indicate a dissociation between neuronal activity and hemodynamics, prompting assessment of NVC in both the time domain (*i.e.*, HRFs) and frequency domain (*i.e.*, TFs). DOI narrowed stimulus-evoked HRFs and enhanced transduction of higher frequency (>0.5 Hz) neuronal activity into hemodynamics. Under resting-state conditions, DOI dramatically altered the HRF shape (*i.e.*, acausal peak), a phenomenon absent in stimulus-based HRFs. These DOI-induced alterations in NVC were region-specific and especially prominent in higher-order brain regions (*e.g.*, retrosplenial cortex). DOI-induced alterations in NVC were reflected in differing accounts of RSFC changes under DOI, as reported by calcium versus hemodynamic activity. Such dissociations carry widespread implications regarding the interpretation of purely hemodynamic measures of brain function, such as human BOLD-fMRI.

### 3.2) Prior research

#### 3.2.1) Neuronal signatures of psychedelic exposure

Classic psychedelics are 5-HT**_2A_** receptor agonists with hallucinogenic properties [10, 32–35, 83–85]. This receptor subtype is predominantly located in cortical regions associated with “higher” cognitive functions (*e.g.*, default mode network, and anterior cingulate cortex) [38, 86–89], regions thought to be key mediators of the psychedelic experience [33, 37, 86–92]. Invasive electrophysiological studies have demonstrated regionally specific modulations of neuronal excitability following psychedelic administration [17, 55–57, 59, 93–95]. For instance, administration of DOI increased pyramidal neuronal firing rate and decreased the low-frequency power in local-field potentials within rat prefrontal cortex. These effects were only partially reversed by MDL, potentially reflecting DOI’s and MDL’s affinity for other receptors such as 5-HT**_2C_** receptors [55, 96]. Although these studies are informative, they reflect regional expression of receptor subtypes [93, 97–99] and may not capture brain-wide emergent phenomenology [100], as captured by network-level neuroimaging techniques like fMRI and WFOI.

#### 3.2.2) Interpretation of psychedelic agents on resting-state fMRI in humans

Recent resting-state fMRI studies in humans have indirectly examined the brain-wide neuronal effects of psychedelic agents [14, 15, 18, 20, 71, 72, 74, 101–105]. Notwithstanding some preprocessing– and experimentally-dependent inconsistencies [99, 106], there is a broad consensus that psilocybin reduces functional connectivity within association networks, particularly within the default mode network (DMN) [14, 15, 74, 81, 105]. This reduction has been hypothesized to underlie the subjective experience of ego dissolution [11, 16, 20, 34]. Many studies also report increased functional connectivity between association networks and other regions of the brain [16–18, 20, 59, 99, 100], suggesting a reorganization of brain-wise communication patterns.

Psychedelics induce dissolution and fragmentation of canonical RSFC patterns, which have been quantitated using entropic measures [11, 18, 44, 72, 100]. This reorganization is reflected in reduced modularity (*i.e.*, decreased intra-network connectivity with enhanced inter-network connectivity) [16, 18, 80–82]. Notably, the DMN – a constellation of brain regions consistently impacted by psychedelics [14, 19, 70, 81, 107] – exhibits both decreased intra-network and increased inter-network connectivity [14, 70, 81, 107]. Together, these findings suggest that psychedelics modulate neural activity throughout the brain. However, interpreting purely hemodynamic measures as indicative of neural activity is inadequate without accounting for the neurovascular effects of psychedelics.

We observed similar alterations in network integration in neuronal reports of brain function. This effect was dampened in hemodynamic reports, demonstrating the potent effects of NVC on neuronal assessment of functional brain organization. Importantly, our findings present a more nuanced picture than previous fMRI reports, which have generally indicated decreased intra-network and increased inter-network connectivity within and between the DMN. Critically, our results highlight differential connectivity properties in the anterior and posterior DMN regions when assessed using neuronal versus hemodynamic estimates of RSFC.

#### 3.2.3) Neurovascular coupling

Apart from neuromodulation [108–113], serotonin — the endogenous ligand for the 5-HT**_2A_**R — also has potent vasoactive effects [36, 37, 114]. For example, 5-HT**_2A_**R binding triggers vasoconstriction through intracellular signaling pathways involving phospholipase C, intracellular calcium release, and activation of kinases such as protein kinase C phosphorylation and CaMKII [115]. Moreover, serotonin can directly modulate blood flow by activating serotonin receptors on blood vessels [114]. Further, 5-HT**_2A_**Rs are expressed on astrocytes [38], cells that play a crucial role in NVC [39–43, 116–118]. Serotonin might also indirectly affect NVC through its ability to modulate systemic physiology [114, 119, 120] (*e.g.*, respiratory rate, cardiac rate, blood pressure), which can influence neurovascular relationships [65, 121–127]. For example, variations in cardiac rate can modulate low frequency BOLD power in a region-specific manner [128], a poorly understood phenomenon owing to the numerous factors that influence fluctuations in arterial pressure and parasympathetic/sympathetic nervous activity [129, 130]. These modulations in physiology can directly affect estimates of hemodynamic-based RSFC when assessed by hemodynamic measures like fMRI, fNIRS, and optical intrinsic signal imaging [65, 121–128, 131–135].

Prior psychedelic research has demonstrated dose-dependent [136–140] effects on systemic physiology, such as increased cardiac rate and blood pressure, further demonstrating the possibility that psychedelic agents modulate NVC and BOLD signaling [34, 141]. However, it is important to note that methylphenidate, the non-psychedelic control condition in human studies, also produces significant physiological effects (*e.g.*, increased blood pressure and heart rate [142]) without altering stimulus-based fMRI HRFs. This crucial control data suggests that the potent psychedelic NVC effects are not solely a result of systemic physiological changes.

### 3.3) Hallucinogenic doses of DOI disrupts neurovascular coupling

DOI increased calcium activity in the delta band. This observation is concordant with prior EEG studies in humans on the effects of other psychedelics [57, 58, 143, 144]. In the present data, increased delta band activity (frequencies currently inaccessible to most fMRI studies) was also observed in hemodynamic signals (*e.g.*, transfer functions and hemodynamic power). Enhanced delta-band transduction and decreased HRF width was particularly pronounced in lateral portions of the mouse cortex. This topographical pattern, which reflects regional differences in NVC, may correspond to a key psychedelic effect in humans: the loss of functional segregation between primary sensory cortical areas (*e.g.*, somatosensory and visual) and the default mode network, observed in the infraslow band frequencies below 0.1 Hz [14, 15, 19, 70, 81, 145].

In contrast to a hallucinogenic DOI dose, we did not observe altered NVC from an empirically derived, sub-hallucinogenic dose of DOI and the non-hallucinogenic, 5-HT**_2A_**R agonist Lisuride (**Fig. S14**). Additionally, the broad scope of 5-HT**_2A_**R function could be due to possible biased agonism [146]; both DOI and Lisuride [147] target the 5-HT**_2A_**R receptor, but activate different signaling cascades once bound [148]. DOI strongly activates the Gq-coupled cascade, leading to the phosphorylation of signaling proteins such as phospholipase C (PLC), protein kinase C (PKC), ERK/MAPK, and CaMKII. This activation increases the production of secondary messengers like inositol phosphate signaling molecules (IP3) and diacylglycerols (DAG), which influence synaptic activity, neuronal excitability, and blood flow regulation. These properties are especially relevant in the context of NVC – increased production of IP3 in astrocytes during neuronal activity triggers a calcium signal that activates pathways leading to the release of epoxyeicosatrienoic acids (EETs) and prostaglandin E2 (PGE2), resulting in nitric oxide release and vasodilation [149]. In contrast, Lisuride’s influence on these pathways is weaker and leads to less phosphorylation of signaling proteins and less production of secondary messengers regulating blood flow [148]. The ability of DOI to engage both the PLC and phospholipase A2 pathways results in notable vascular and neural effects, while Lisuride’s activation at the 2A receptor leads to different engagement of downstream pathways presumably responsible for hallucination.

Furthermore, within the central nervous system, 5-HT2**_A_**R are expressed not only in neurons but also astrocytes [38] and other cell types that modulate NVC [39–43]. The broad action of DOI across many brain regions and receptors [150–154] indicates that this ligand is not purely selective for 5-HT**_2A_**Rs but can also bind to many other 5-HTRs and Class I G protein-coupled receptors. Direct action on 5-HT2**_A_**R-expressing neurons *via* DOI, or neural activity downstream of 5-HT2**_A_**R neurons, might also contribute to neuromodulatory effects that influence hemodynamic activity. Given inability of DOI to drive pyramidal neurons to fire in slice preparations (**Extended Data Fig. 3**), we suspect that most calcium effects observed in WFOI arise from such downstream neural activity.

#### 3.3.1) A potential retrograde calcium-to-hemodynamic relation

DOI induced an apparent zero-lag notch in the HRFs resulting in a peak at negative lags. These observations potentially imply a relation in which hemodynamic activity precedes calcium activity. Importantly, the prominence of this effect was regionally dependent (*e.g.*, largest in secondary motor and retrosplenial regions). Furthermore, this feature was present in calculations of lagged cross-covariance between unfiltered calcium and hemodynamic signals and appears in the frequency domain as a broad peak around ∼0.8 Hz in the same regions. A constant slope in the phase plot indicates a fixed temporal lag [155]. On average, before compound injection, the calcium-to-hemodynamic phase relation exhibited a positive slope at frequencies between zero and ∼1 Hz, indicating that calcium leads hemodynamics activity in this band. However, DOI (and sometimes DOI+MDL) induced negative phase slopes in specific cortical regions within the same frequency band (*e.g.*, frontal, and cingulate regions). This result suggests that hemodynamics lead calcium in these regions, a remarkable difference from the baseline state.

The presence of the notch and peak at negative lags could potentially arise from DOI-induced alterations in the calcium-hemodynamic relation (a neurobiological account) or misinterpretation of the optical data (artifact). In the artifactual account, the notch may represent the superposition of high-frequency anticorrelations on a temporally broad lag-dependent relation. Given that this notch was unevenly distributed across the cortex, only appeared under DOI, and was absent in stimulus-evoked HRFs subjected to the same analysis, we believe it is highly unlikely to be an artifact.

An alternative account of this feature includes two, DOI-dependent, neurobiological phenomena: neuro-vascular coupling and vasculo-neural coupling (VNC) [64, 127, 156–158]. Many mechanisms that induce functional hyperemia often impact neuronal activity. Chemical factors that modulate blood flow often also regulate neuronal function [157]. For example, freely diffusing nitric oxide (NO) causes relaxation of smooth muscle [159] and modulates neuronal activity [160]. Blood volume changes during functional hyperemia, can also lead to morphological changes in surrounding neurons, activating mechano-sensitive ion channels (*e.g.*, amiloride-sensitive Na+ channels and stretch-activated cation channels, both of which are abundant in the mouse neocortex [161]). Additionally, brain temperature, which is directly governed by blood flow [162], can affect neuronal activity; a temperature decrease of 2 degrees Celsius is sufficient to reduce neuronal firing rates [163]. Furthermore, vaso-astrocytic interactions can indirectly impact neuronal activity through diffusible messengers (*e.g.*, direct contact between astrocytic end feet and capillary endothelial cells [164] can modulate endothelial NO release triggered by increased hemodynamic activity [165–167]) and mechanical interactions (*e.g.*, stretch-activated ion channels in astrocytes [168]). Furthermore, activation of 5-HT receptors themselves implicate many possible VNC mechanisms. For instance, 5-HT activity increases astrocytic calcium activity [169], modulates NO activity [170–172] and decreases brain temperature [173]. Considered collectively, there is ample evidence that psychedelics could give rise to VNC both directly and indirectly.

### 3.4) Limitations

It is important to note the changes in NVC we observe are specifically between Thy1-expressing cortical neurons (primarily expressed in pyramidal cells of cortical layers 2/3 and 5) and the vasculature. Several studies, including our own, have demonstrated strong spatiotemporal coupling between Thy1-based activity and subsequent hemodynamic activity under both task and resting-state conditions [51, 174–178]. However, activity in subpopulations of excitatory and inhibitory neurons can be sufficient to alter cerebral blood flow [112, 177–184]. Furthermore, hemodynamic responses can vary depending on which cell populations are activated, sometimes even exhibiting opposing or biphasic responses [184–188]. These vasoactive effects are especially significant in the context of psychedelics, as 5-HT**_2A_** receptors are predominantly expressed in pyramidal cells—particularly in the prefrontal cortex—and, to a lesser extent, in certain subsets of interneurons [87, 189]. Within the cortex [92, 190] and deeper brain structures [87, 191–193], serotonergic modulation of inhibitory-excitatory balances [194–196] lead to profound effect on local and global regulation of brain perfusion.

Determining the degree to which 5-HT**_2A_**R agonism, or psychedelics more broadly, affects other cell type-specific coupling relationships with the vasculature will help elucidate mechanisms underlying altered hemodynamic signaling. Future research could incorporate electrophysiological measurements to determine the local contributions of excitatory and inhibitory neurons (*i.e.*, comparing excitatory activity to total activity). Alternatively, a more global characterization of neurovascular relationships could be achieved by using mice expressing genetically encoded calcium or voltage indicators in other neurons [197–202] or glia [203, 204].

Several observations under DOI conditions were not fully reversed by the 5-HT**_2A_**R antagonist, MDL, which raises the question of receptor-dependent psychedelic effects outside of 5-HT**_2A_**R binding [150–154]. MDL is one of the most selective 5-HT2**_A_**R antagonist currently available. Although it retains some affinity for 5-HT2CRs, sigma, α_1A_, and α_2B_ receptors, there is limited *in vivo* evidence suggesting it acts strongly at these receptors at the dose used in this study [205]. Similarly, Lisuride, the putatively non-hallucinogenic, 5-HT2AR agonist, has a different affinity profile for these and other 5-HT receptors compared to DOI. However, exposure to Lisuride did not alter neurovascular coupling or induce head twitch responses in mice. Investigating a broader range of compounds [147, 206, 207] and receptor subtypes would provide a more comprehensive understanding of the neurophysiology underlying psychedelic effects.

Although our NVC model accounts for many features of the observed data, it assumes causality, linearity, and time-invariance. Collectively, these assumptions may oversimplify the complexities of NVC. The most salient example of this oversimplification is the HRF measured after DOI, which contains a primary (positive lag) and secondary (negative lag) peak, challenging the assumption of causality. Therefore, more comprehensive models of coupling between neuronal and hemodynamic activity could account for intermediate steps [208–211] or more complex retrograde phenomena (*e.g.*, VNC) [64, 127, 156–158]. Expanding our NVC model to account for nonlinear, acausal, self-modulation/autoregressive, or cross-frequency coupling effects [135, 212–214] could characterize other aspects of psychedelic-induced NVC alterations.

Emerging evidence also suggests sexual dichotomy in the pharmacokinetic and behavioral profile of multiple psychedelics [215–218]. Although this study included both sexes, it was not adequately powered to evaluate sex-specific differences in outcome. Future studies will be critical to evaluate the degree to which different psychedelics affect sex-specific alterations in neurophysiology.

Lastly, saline often altered neurophysiology (excluding NVC), predominantly in a frequency-dependent manner. This phenomenon was also observed in the independent dataset examining Lisuride (**Extended Data Fig. 9f**). These changes may be attributed to the mouse experiencing increased stress after injection due to handling and needle insertion. Such effects can manifest as decreased delta-band power [219, 220]. To that end, assessment of other compound conditions are interpreted with respect to these and other inherent “stress-inducing” effects.

### 3.5) Conclusions and next steps

Our findings raise questions that can only be addressed with more direct measures of neuronal activity and NVC. In human studies, this could be achieved with simultaneous EEG-fMRI [221–223] and consideration of NVC. Our WFOI sampling rate resulted in sensitivity to hemodynamic activity in frequencies extending up to ∼10 Hz. The largest hemodynamic effect of DOI occurred at frequencies above 0.2 Hz. The extent to which this phenomenon is relevant to human fMRI studies is not clear as BOLD-fMRI signals are markedly attenuated at frequencies above ∼0.2 Hz in awake humans [224, 225]. To address this limitation, optical techniques such as diffuse optical tomography could be used to measure cortical hemodynamics with higher temporal resolution [226].

More broadly, psychedelic research could benefit from a more cautious stance towards the conventional neurocentric interpretation of BOLD-fMRI [227]. This shift encourages a broader, more holistic perspective concerning the genesis of blood-based signals [39–43, 116–118, 121–123, 127, 228–230]. Such an approach could improve our understanding of the effects of psychedelics on brain function.

## 4) Methods

### 4.1) Analysis of hemodynamic response functions in task-based human fMRI data

We assessed changes in the hemodynamic response functions (HRFs) of previously published psychedelic human fMRI task data (**Figs. 1a** and **Extended Data Fig. 1a**) [44]. Briefly, subjects were imaged under no compound, methylphenidate (40mg), and psilocybin (25mg) while performing a simple auditory-visual matching task. Generalized linear models (GLMs) were computed at pre-specified regions of interest (ROIs) which were selected from the Gordon-Laumann parcellation [231] (left/right calcarine sulcus (V1), left/right auditory cortex (A1), left language area (Wernicke-like area), left hand knob, left angular gyrus, and right angular gyrus (default mode). HRFs were fit using a double Gamma function, which can be described using thee parameters (peak value, dispersion, and time-to-peak) [232]. MANOVAs were used to assess the differential effect of compound (or no compound) and participant on each parameter. A post-hoc t-test was conducted on regions where the MANOVA indicated a significant effect of compound.

### 4.2) Mice and animal preparation

All mice were maintained in the animal facilities at Washington University in Saint Louis and all procedures were performed in strict accordance with Washington University in Saint Louis Animal Care and Use Committee requirements. Eight transgenic mice (4 males, 4 females) expressing jRGECO1a driven by the thymus cell antigen 1 (*Thy1*) promotor were used (JAX Strain: Tg(Thy1-jRGECO1a) GP8.20Dkim/J; stock: 030525) [233–235] to visualize cortical calcium fluctuations primarily from excitatory neurons of layers 2/3 and 5 in relation to hemodynamic activity [235, 236].

Prior to imaging, a cranial window was secured to the intact skull of each mouse using dental cement, following scalp retraction under 2% isoflurane anesthesia, as described in our previously published protocols [237] (**Fig. 1b**). One hour before surgery, mice were given buprenorphine-SR (1.0 mg/kg subcutaneous) and recovered from surgery for three days prior to any handling or experimentation. Once recovered, and prior to any imaging experiments, all mice were behaviorally acclimated to the awake imaging apparatus for 45 minutes per day for 7 days to reduces stress as per our previously published protocols [45, 238].

An initial pilot study (4 females, 5 males) using saline and Lisuride [147, 206] under awake, resting-state conditions was conducted to determine the appropriate number of subjects and experimental design. Mice were imaged for 30 minutes before injection and for 90 minutes immediately following the injection of Lisuride. Because we observed the largest effects of Lisuride on cortical activity (see **Extended Data Fig. 9f**) 30 to 60 minutes post-injection, all subsequent WFOI occur before and 30-60min after injection of any compound. These results are in line with prior work using DOI demonstrating the largest effects occur 30 minutes after injection [239].

Compound dosages were determined from prior literature and through evaluation of behavior responses, and particularly the head twitch response (below) [90, 240–243]. From these pilot studies the following dosages were used for experimentation: DOI (4 mg/kg), MDL (0.10 mg/kg), DOI+MDL, and saline. To assess the effect of sub-hallucinogenic doses and non-hallucinogenic 5-HT**_2A_**R agonists on NVC, low-dose DOI (0.04 mg/kg, as determined HTR) and Lisuride (1.0 mg/kg) were also used. To control for volumetric effects, all mice were injected with the same volume of liquid per mass (20 mL/kg). All compounds were injected into the intraperitoneal (i.p.) space and prepared on the day of experimentation.

### 4.3) Head twitch response and dose response

The head-twitch response (HTR) is a widely used behavioral assay in which rodents exhibit rapid side-to-side head movements after administration of serotonergic hallucinogens and other 5-HT**_2A_**R agonists. HTRs strongly correlate with the duration of action and strength of these compounds [90, 240–243]. Extensive research has demonstrated that DOI induces powerful psychedelic effects and induces HTRs [90, 241, 244, 245]. A custom magnetometer was built for automating HTR characterization based on open-source designs [240]. Briefly, a Plexiglas chamber (4” H x 4” W x 4” L) was constructed and wrapped with 820 feet of 30 AWG enameled copper wire to create a solenoid with 154 turns. The solenoid output was connected to an amplifier (Phono preamp; PP444, Pyle). Magnetic tags were glued on the mouse ear (3mm diameter, 0.5mm thickness, ∼58 mg) so that when the mouse was placed in the chamber, any head twitches would induce an electromotive force in the coil (**Fig. S1**).

Raw HTR data were denoised using a discrete wavelet transform (wavedec, MATLAB) with a symmetric wavelet of order 6 (i.e., ‘sym6’ MATLAB). Soft thresholding was performed using Stein’s unbiased risk estimation for each level’s detail coefficients before reconstructing the signal. Once denoised, processing of HTR time courses followed prior work [240].

Pilot studies in 3 mice and 4 conditions (saline, DOI: 0.04 mg/kg, DOI: 0.4 mg/kg, and DOI: 4mg/kg) were used to validate our magnetometer and automated detection algorithm by comparing against results from a manual HTR scorer blinded to condition (R^2^=0.950 between automated and manual scoring of HTRs; **Fig.S2a**). These data were also used to assess the dose response of DOI (**Fig. S1b**). A second cohort of mice (2M, 2F) was used to assess HTRs following randomized injection of each compound (Saline, DOI, MDL, DOI+MDL; same dose and concentration as WFOI experiments; **Fig.S1c**). Lastly, a third cohort of mice (2M, 2F) was used to assess HTRs following of Saline, Lisuride (1.0 mg/kg), and a sub-hallucinogenic (low dose) of DOI (0.04 mg/kg, as determined by compound response curve) and underwent the same protocol as cohort 2. For all cohorts, HTR recordings began 30 minutes after injection of compound. For cohort 1 (HTR dose response), recordings were 10 minutes long, and for cohorts 2 (main manuscript compounds) and 3 (low dose DOI and the non-hallucinogenic, 5-HT2**_A_**R agonist, Lisuride), recordings were 30 minutes long (**Extended Data** Fig 9a). For all cohorts, tests of compound effects on individual mice were separated by at least 3 days to avoid potential changes in tolerance.

### 4.4) Ex vivo electrophysiological and imaging recording

Adult Thy1-RGECO1a (n= 5) and C57BL/6J (n = 3) mice were anaesthetized *via* i.p. injection of a cocktail containing ketamine, xylazine and acepromazine (69.57 mg/ml; 4.35mg/ml; 0.87mg/ml; i.p. 182mg/kg, respectively). Mice were then perfused with iced cold slicing-aCSF consisting of 184 mM sucrose, 2.5 mM KCl, 1.25 mM NaH**_2_**PO**_4_**, 10 mM MgSO**_4_**, 20 mM HEPES, 30 mM NaHCO**_3_**, 25 mM glucose, 0.5 mM CaCl**_2_**, 5 mM sodium ascorbate and 3 mM sodium pyruvate, oxygenated with 95% O**_2_** and 5% CO**_2_**. The pH was 7.3–7.4 and osmolality adjusted to 315–320 mOsm with sucrose. The forebrain was dissected and embedded with 2% agarose in slice-aCSF and coronal slices were cut with 300 μm thickness using a vibratome (VF310-0Z, Precisionary Instruments, MA, USA). Slices were incubated at elevated temperature (32°C) for 30 minutes and then transferred to room temperature in holding-aCSF consisting of 92 mM NaCl, 2.5 mM KCl, 1.25 mM NaH**_2_**PO4, 30 mM NaHCO**_3_**, 20 mM HEPES, 25 mM glucose, 2 mM MgSO**_4_**, 2 mM CaCl**_2_**, 5 mM sodium ascorbate and 3 mM sodium pyruvate, oxygenated with 95% O2 and 5% CO2. pH was 7.3–7.4 and osmolality adjusted to 310–315 mOsm. The slice preparation procedures were modified from previous study [246]. Both electrophysiological and imaging recordings took place in a recording chamber mounted on upright microscope (BX51WI, Olympus Optical Co., Ltd, Tokyo, Japan) as previously described [247] with continuous perfusion of warm (29–31°C) recording-aCSF consisting of 124 mM NaCl, 2.5 mM KCl, 1.25 mM NaH**_2_**PO**_4_**, 24 mM NaHCO**_3_**, 5 mM HEPES, 12.5 mM glucose, 2 mM MgCl**_2_**, 2 mM CaCl**_2_**, oxygenated with 95% O2 and 5% CO2 and pH 7.3–7.4 with osmolality adjusted to 305–310 mOsm using sucrose. Cell-attached recordings were performed through borosilicate glass pipette (GC150F-10, Warner Instruments, Hamden, CT, USA) with a resistance around 6-9 MΩ when filled with recording-aCSF. Signals were gathered using Multiclamp 700B amplifier (Molecular Devices, San Jose, CA, USA) and digitized at 10k Hz via Axon Digidata 1550B interface (Molecular Devices, CA, USA) coupled with Clampex software (Molecular Devices, CA, USA). A subset of cell-attached recordings were performed in C57BL/6J mice without calcium imaging due to mouse availability. Calcium imaging in Thy1-RGECO1a recordings were collected by epifluorescence equipment mounted on microscope coupled with a highspeed camera (ORCA-Flash4.0LT, Hamamatsu Photonics, Shizuoka, Japan).

### 4.5) Imaging Hardware

The experimental setup included three separate cameras (**Fig. 1c**), one for optical neuroimaging (WFOI) and two for movement monitoring (one for monitoring murine motion, and one for monitoring pupil dynamics).

#### 4.5.1) Wide-field Optical Imaging (WFOI)

Awake cortical neuroimaging was performed using a modified version of a previously-published WFOI system [63]. A custom light engine consisting of 470nm (measured peak λ = 457 nm, LCS-0470-50-22, Mightex Systems), 530nm (measured peak λ = 525nm, LCS-0525-60-22, Mightex Systems), and 625nm (measured peak λ = 637 nm, M625L3, Thorlabs) light-emitting diodes (LED) illuminated the skull (**Fig. 1c**). Diffuse reflected light for optical intrinsic signal imaging and fluorescence emission were collected by a lens (focal length = 75mm, NMV-75M1, Navitar), split by a 580nm dichroic (FF580-FDi02-t3-25×36, Semrock) and sampled by two scientific complementary metal–oxide–semiconductor (CMOS) cameras with USB3 connectivity (Zyla 5.5, Andor). A 500nm long pass filter (FF01-500/LP-25, Semrock) in front of Camera 1 (CMOS1) passed 530nm reflectance. The 580nm dichroic and a 593nm long pass filter (FF01-593/LP-25, Semrock) in front of Camera 2 (CMOS2) blocked 530nm jRGECO1a excitation light and passed jRGECO1a emission and 625nm reflectance. A separate 700 nm short pass filter (Thorlabs, FESH0700) placed behind 593nm long pass filter and in front of CMOS2 blocked near-infrared illumination used for movement monitoring (see below). Each WFOI camera sensor was cropped to 1024×1024 pixels, and 2×2 binning was performed to increase the acquisition rate and improve the signal-to-noise ratio (SNR) of each image. The frame rate of each camera was 60 Hz, with all contrasts imaged at 20 Hz. The LED and camera exposures were synchronized and triggered *via* a data acquisition card (PCI-6733, National Instruments) using MATLAB R2022 (MathWorks). LED spectra were measured using an Ocean Optics USB 2000+ spectrometer.

#### 4.5.2) Motion and Pupil Tracking

Mouse motion and fluctuations in pupil diameter were each tracked by CMOS cameras (Thorlabs, DCC3260C) placed in front of the mouse (see below, **Extended Data Fig. 2** and **Fig. S3**). The motion camera was placed approximately 6 cm away from the mouse’s snout and captured spontaneous facial and forepaw movements. The pupil camera was places approximately 3 cm away from the mouse head. The fields-of-view of each camera were as follows: Motion FOV: ∼70 mm-by-70 mm, 480 pixels-by-480 pixels, 0.146 mm/pixel; Pupil FOV, ∼8 mm-by-8mm, 480 pixels-by-480 pixels, 0.017 mm/pixel). Both FOVs were illuminated by a 780 nm LED (Thorlabs, M780L3). The frame rate of each camera was 20Hz and was time-locked to the 625nm LED used for WFOI. Specular reflection in Motion/Pupil image sequences was minimized using crossed near-infrared, linear polarizers (Thorlabs, LPVISC100).

### 4.6) Data Acquisition

During imaging, mice were supported by a felt pouch and mounted to the WFOI system using an aluminum bracket attached to the cranial window [45, 63, 184, 238, 248]. The awake neuroimaging paradigm consisted of interleaved epochs of “resting-state” and “stimulus-evoked” imaging, as performed with human fMRI [249] (**Fig. 1d**, top). “Resting-state” is experimentally defined as the subject in the recording apparatus, isolated from extraneous stimuli and not subjected to any experimentally-imposed stimuli (*e.g.*, overt sensory inputs) [54]. While the animals are head fixed and secured in the felt pouch (which limits any head motion and large movements of the limbs) they are free to look around, shift their body position, whisk, groom, *etc*. Resting-state epochs were 180 seconds long. Stimulus-evoked responses were elicited by delivering puffs of air (40 PSI, 100 ms pulses at a frequency of 3 Hz) to the right whiskers for 10 seconds, followed by a 20-second recovery period. This resulted in a total stimulus block length of 30 seconds (**Fig. 1d**, bottom). Before and after injection, resting-state and stimulus-evoked epochs were each repeated nine times, resulting in 27 minutes of resting-state data and 4.5 minutes of task data, for a total of 31.5 minutes. Our initial pilot studies determined the largest

### 4.7) Motion and Pupil Tracking

Prior to any analysis, each pixel’s time trace was normalized by its temporal mean (*via* division) to account for illumination inhomogeneities. Motion image sequences were spatially decimated by a factor of two to expedite processing. Optical flow (OF) was performed on all motion monitoring movies, as previously reported [250] (**Extended Data Fig. 2a**,i**-iii**). OF estimates were generated using the Lucas-Kanade method [251, 252]. This algorithm produces a spatiotemporal series of vectors where, for each frame, a pixel’s vector represents the change in direction and magnitude over the next frames (n = 5; **Extended Data Fig. 2a,iv**). The root-sum square (RSS) of each frame’s vector magnitude field was computed to yield a single time-trace estimating movement (large values signify large changes in movement; **Extended Data Fig. 2a**,v).

Movement was also assessed using a different algorithm to demonstrate robustness of movement analysis to choice of algorithm. Specifically, the temporal derivative of each pixel’s time trace (from spatially decimated movies) was computed to estimate pixel-wise motion (**Fig. S3a**). Histograms of these motion-characterizing time traces were calculated and averaged across mice before and after compound injection (**Fig. S3b**). Spectral content of RSS movement traces were calculated using a smoothed power spectrum density estimate (pwelch, MATLAB 2022). Band-limited integrated power was calculated over the infraslow band (ISA, 0.02 – 0.08 Hz), delta band (delta, 0.5 Hz – 4.0 Hz), and the frequency range between these bands (intermediate, 0.08 Hz – 0.5 Hz; **Fig. S3c**).

Additionally, pupil dynamics are associated with spontaneous changes in arousal, which give rise to spontaneous behaviors such as whisking, grooming, ad fidgeting [135, 253–255]. Pupil dynamics were therefore monitored and tracked (*via*, an in-house, deep-learning segmentation algorithm) and compared across compounds (**Extended Data Fig. 2d-g**). Specifically, changes in pupil area were synchronized with the motion monitoring and WFOI cameras to determine whether compounds altered pupil dynamics during neuroimaging sessions. To extract pupil area from each image sequence, transfer learning from the pre-trained Segment Anything Model (SAM) [256] was employed utilizing the Vision Transformer model [257] (*i.e.*, ‘facebook/sam-vit-huge’) in the SamProcessor framework. This model, initially trained on a large dataset, underwent fine-tuning to specialize in accurately delineating pupil contours for each frame. To optimize the model’s performance for the specific task of pupil segmentation and ensure effective knowledge transfer, all encoder layers were frozen. This strategic freezing of layers enabled the model to retrain its learned representations of general features while allowing the trainable parameters of the decoder to adapt specifically to our task (**Extended Data Fig. 2c**).

Adam optimization with an initial learning rate of 1E-3 and a scheduler that reduces the learning rate by 25% every 50 epochs was employed for training. Monai’s DiceFocalLoss (gamma value of 2.0) was utilized instead of the more conventional Dice or Dice cross-entropy losses due to the substantial class imbalance within the imaging frames (>80% of pixels were non-pupil). To mitigate overfitting, a dropout rate of 30% and a regularization constant of 1E-4 were applied. Training spanned 100 epochs with early stopping implemented to avoid overfitting—training stopped if validation loss at epoch n was less than the median validation loss between epochs n-15 to n.

For the first round of training, all pupil videos were clustered (k-means) with 80 clusters determined via elbow plot. The cluster centroids representing the largest clusters (the 10 clusters with the most data points) were manually segmented. At least 15 user-defined points were used to contour the pupil, which was then fit to an ellipse using a least-squares approach (**Extended Data Fig. 2d**). This process resulted in the labeling of 956 images. Data augmentation techniques were applied, including affine transformations with randomly selected parameters and random horizontal flips. Consequently, the final dataset comprised 1912 pupil images along with their respective labels. Of these, 80% were allocated for training and 20% for validation.

In the subsequent training phase, 15 pupil videos were randomly selected for frame-by-frame processing using the trained model. Poorly segmented frames often resulted in a large spatial shift in the likelihood distribution/map. (**Extended Data Fig. 2e**). Therefore, for each frame, the center of mass (COM) of the likelihood map was calculated and, for each video, the ten largest deviations of COM were identified. Frames before, at, and after these deviations were manually segmented (15 images per video). This process yielded approximately 450 additional labeled frames, which were also subjected to augmentation. These frames were appended into the training dataset, and the training process (i.e., starting with the SAM initial weights), along with outlier detection and refinement, was repeated once more. In this way, the training dataset was built and fine-tuned to model weaknesses before the final training.

Once the model was adequately trained, every pupil video was analyzed at its native frame rate (i.e., 20 Hz). The sequence of spatiotemporal likelihood maps was then median filtered using a 5-frame kernel (R2R Fig. 1.6d). For each frame, contours (imcontour and roipoly, MATLAB) were fit around the likelihood cloud and all pixels within this contour were counted, resulting in a time course of pupil area changes during imaging (example of pupil trace in **Extended Data Fig. 2f**). The temporal derivative of each pupil area time courses was evaluated for each mouse and compound condition to determine whether changes in pupil diameter different pre and post injection. Because mice often closed their eyes during whisker stimulation, pupil analysis was only performed during resting-state epochs.

### 4.8) WFOI data pre-processing

#### 4.8.1) Spatial normalization

Image sequences from each neuroimaging camera were spatially coregistered using a projective transform, which was constructed from four user-selected landmarks present within both camera’s FOV. WFOI image sequences were then coregistered to the Allen Mouse Brain Atlas following our prior work [61, 237]. Briefly, two anatomical landmarks are identified and labeled on the cortex (1) the junction between the rostral rhinal sinus and the sagittal sinus and (2) lambda. All pixels labeled as brain were used to create a brain mask for each mouse. All subsequent analyses are performed within this common space.

#### 4.8.2) Spectroscopy and spatiotemporal filtering

For each WFOI camera, one second of dark frames were collected, averaged, and subtracted from image sequences [237]. Raw WFOI image sequences (512×512 pixels) were spatially binned (4×4) off camera to further increase the SNR of each image, resulting in 128×128 pixel images. For each channel, slow trends in light levels were temporally detrended with a 5^th^ order polynomial fit [45, 61, 79, 237]. Images were spatially smoothed using a Gaussian filter (5×5-pixel kernel, standard deviation of 1.2 pixels). Changes in wavelength-dependent diffuse reflectance were converted to changes in hemoglobin concentration using the modified Beer-Lambert law [61, 237]. Specifically, for each pixel, ϕ_λ_(t) = ϕ_0,λ_exp (−Δμ_a,λ_(t)X_λ_), where ϕ_λ_(t) is the measured light intensity at each wavelength, ϕ_0,λ_ is the average intensity over the imaging session, μ_a,λ_ is the change in absorption coefficient due to changes in blood volume, and X_λ_ is the differential pathlength that accounts for the volume of tissue sampled by photons at each wavelength. Isolating Δμ_a_ yields 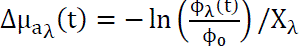, where λ indexes the channel used (*e.g.*, LED centered at 530 nm). The system of equations Δμ_a,λ_ = E_λ,i_Δ[Hb_i_](t) was solved, such that E is the wavelength-dependent extinction coefficient for the i^th^ hemoglobin species (*i.e.*, oxy– and deoxy-hemoglobin). Hemoglobin extinction coefficients were those tabulated by Scott Prahl [258]. Differential pathlengths were computed assuming a semi-infinite geometry, initial concentration of hemoglobin (sO_2_ = 71%, [HbT]_initial_ = 76 micromol), and estimates of the reduced scattering coefficient [259]. Calculation of extinction coefficients and differential path lengths were weighted by LED spectra. All optical properties are reported in **Table S1**. All hemodynamic outcome measures were reported using total hemoglobin (HbT).

To accurately assess fluorescent emission in media with changing optical properties, such as varying chromophore concentrations, it is essential to correct for absorption artifacts. Accordingly, raw jRGECO1a fluorescence was corrected using estimated changes in Δμ_*a*_ at excitation and emission wavelengths, as established in prior work [50–52]. Corrected temporal fluctuations in fluorescence at each pixel were converted to percent change: 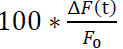, where F(*t*) is the corrected fluorescence, F0 is the temporal mean of F, and ΔF(t) = F(t) − F_0_. The efficacy of the hemodynamic correction was determined in mice (2M, 1F) expressing the yellow fluorescent protein (YFP) under a Thy1-promotor, which provides non-activity-dependent fluorescence (**Fig. S2**).

### 4.9) Signal pre-processing

Resting-state, spontaneous fluctuations in calcium and hemodynamic signaling (sampled at 20Hz) were filtered to obtain frequencies between 0.02 Hz – 0.08 Hz to assess infraslow activity, 0.08 Hz – 0.5 Hz to assess intermediate activity, and 0.5 Hz-4.0 Hz to assess delta band activity. Filtered signals were downsampled to 1.25 times the lowpass filter frequency cutoff (*e.g.*, data filtered between 0.5Hz-4Hz were resampled to 5 Hz). For stimulus evoked activity and NVC assessment (see below), data were filtered to retain frequencies between 0.01 Hz and 5 Hz and then downsampled to a sampling rate of 12 Hz. All filtering and downsampling was performed using a 5^th^ order Butterworth filter (butter, MATLAB 2022) and anti-aliasing interpolation (resample, MATLAB 2022). To minimize boundary effects from dividing the data into task and rest epochs, filtering was conducted prior to splicing. After filtering and downsampling, resting-state and stimulus-evoked activity were segmented into periods of rest and task using a Tukey window (with a Tukey parameter of 0.3, which attenuated 15% of the data at each end of the window).

### 4.10) Analysis of Stimulus-evoked Activity

To enhance the spatial specificity of evoked cortical topography, global signal regression (GSR) was applied [260]. To facilitate averaging across task presentations, 5 seconds of activity occurring prior to stimulus onset was averaged and subtracted from each image in the block. Blocks were then averaged together for each mouse and group. Response maps of evoked calcium and hemodynamic activity were created by averaging images between 0.2-5.2 and 2.0-7.0 seconds after stimulus onset, respectively.

To account for potential changes in pre– and intra-stimulus variance, effect sizes of evoked activity (**Fig. 2a**) were quantified using Cohen’s D:

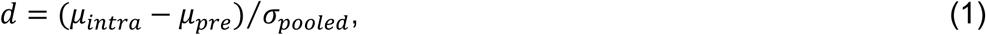

where μ*_pre_*and μ*_intra_* are the mean of the baseline (5 seconds before stimulus onset) and the response period (0.2-5.2 seconds for calcium and 2.0-7.0 seconds for total hemoglobin), respectively, and

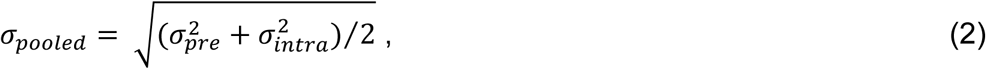

where σ*_pre_* and σ*_intra_* are the temporal standard deviations in activity and *n* is the number of sample points.

To create a high-SNR image representing evoked activity across all mice and groups, all 32 pre-injection effect maps (8 mice·4 compound conditions) were averaged together. From this map, a region of interest (ROI) was defined as all pixels within 25% of the maximum effect (**Extended Data Fig. 1b**). Within the ROI, compound-induced changes in response were assessed using the maximum response amplitude. To account for variability before and during stimulus presentations we also examined maximum response effect. Similarly, the response area was computed using the number of pixels exceeding 75% of the maximum amplitude or the number of pixels exceeding 75% of the maximum effect. The topographical correspondence between evoked calcium and hemodynamic activity was assessed *via* spatial similarities (Pearson correlation) between calcium and hemodynamic effect maps.

Response temporal dynamics for each mouse were evaluated by averaging all pixel time courses within the ROI. Only evoked activity exceeding 2 standard deviations of baseline (pre-stimulus) activity was included in subsequent analysis. Response dynamics are visualized in **Figure 2b**. Changes in evoked activity was compared across groups through quantification of the peak response (the sum between 0.2-2 seconds and 2.2-4.2 seconds, for calcium and HbT, respectively), transient response (the sum between 0-5 seconds), sustained response (the sum between 5-10 seconds), and the post-stimulus undershoot (the sum after stimulus-offset, 10-30 second; **Extended Data Fig. 1c**). Additionally, the effect size between pre-(2-seconds before stimulus-onset) and post-stimulus (0.2-2 seconds and 2.2-4.2 seconds, for calcium and HbT, respectively) was computed as Cohen’s D (**Extended Data Fig. 1c**).

### 4.11) Stimulus-based Neurovascular coupling

Neurovascular coupling (NVC) was modeled as a causal, linear, time-shift-invariant process. For each pixel, let *y* = *X*ℎ where *y* is measured hemodynamic activity, ℎ is the hemodynamic response function (HRF), and *X* is the convolution operator (each column is an *m*-shifted calcium signal). For task, *x* and *y* are the average response from all pixels within the ROI seen in Fig. 2b. In expanded form, this equation becomes:

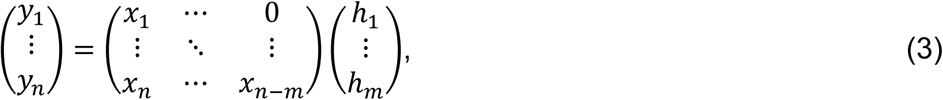

where *n* is the number of sample points and *m* is the length of the convolution kernel. Weighted least squares deconvolution was employed o estimate the HRF:

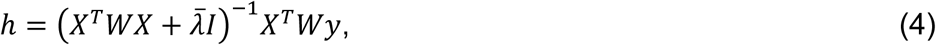

where *W* is a weight matrix, 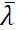 is a regularization constant, and *I* is the identity matrix. We chose the weight matrix to be the singular value spectrum of *X*, namely *W* (*i.e.*, *X* = *U*W*V*^*T*^). To standardize the regularization constant for all mice and conditions, 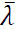 was defined as 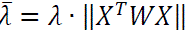, where the fractional regularization constant, λ, was empirically set to λ = 10^−5^.

Prior to deconvolution, an artificial 3 second delay was introduced between and to examine temporal relations prior to time zero (stimulus-based NVC, **Fig. 3**; resting-state NVC, **Fig. 5**). To assess model fit, measured calcium activity was convolved with the HRF to produce predicted hemodynamic signals, which were then correlated with measured hemodynamics (**Fig. S5**). HRFs were parameterized by peak-value, time-to-peak, and full-width-at-half-maximum (**Fig. 3a**, left). Additionally, the oxy-, deoxy-, and total-hemoglobin post-stimulus HRF undershoot (summed activity 2-seconds after stimulus-onset) was compared across compounds (**Fig. S4b**). To assess the frequency-dependent properties of calcium-to-hemodynamic transduction, transfer functions (TF) were defined as the power spectral density estimate of the HRF. Transduction was quantified using integrated power in the ISA, intermediate, and Delta bands (defined above).

### 4.12) Quantification of Resting-state Activity

Resting-state activity was quantified regionally by averaging all time courses across anatomical regions as defined in the Allen atlas [261] (**Figs. 4**, **Extended Data Fig. 4a-b**, and **Extended Data Fig. 5**; Lisuride, **Extended Data Fig. 9f**), which we will refer to as regional signals. Power spectral density estimates (PSDEs) for each region and epoch (9 resting-state epochs, 180 seconds each) were computed using a smoothed spectral estimator (pwelch, MATLAB 2022). These estimates were then averaged across epochs to produce mouse-level PSDEs. Spectra were integrated over the ISA, intermediate, and delta bands (defined above). Global PSDs for each mouse were obtained by averaging the PSDEs across all cortical regions. Finally, group-level cortical resting-state PSDEs were calculated by averaging across all mice.

### 4.13) Resting-state Neurovascular coupling

Resting-state NVC (**Figs. 5**, **Extended Data Fig. 4c**, and **Extended Data Fig. 6**; low-dose DOI, **Extended Data Fig. 9b-c, and Lisuride Extended Data Fig. 9d-f**) was quantified regionally using regional signals as defined in 2.9. Regional time courses were segmented into 30 second, non-overlapping, windows (Tukey window, lobe parameter of 0.3). HRFs were estimated for each 30 second epoch and the median across all epochs defined a regional HRF. Regional estimated HRFs were averaged across regions to obtain a cortical-wide (global) estimate of NVC. HRFs and TFs were parametrized in the same manner as stimulus-evoked NVC.

To better understand emergent features in resting-state NVC process (*e.g.*, peaks at times prior to zero), cross-covariance, coherence, and phase was also computed on the same resting-state epochs (**Extended Data Figs. 7-8**, see below).

### 4.14) Cross-covariance analysis

Regional cross-covariance functions between unfiltered calcium and HbT signals were computed to validate and assess emergent features present in neurovascular coupling estimates obtained using weighted deconvolution (see main text). From these functions, peak and time to peak were calculated for each region. The frequency-dependent content of cross-covariance functions were estimated by as the cross-spectral density, quantified as magnitude-squared coherence (MSC):

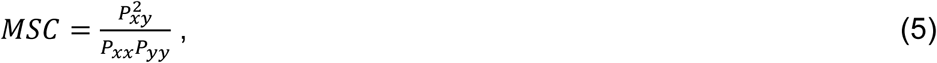

where P*_xy_*, P*_x_*, and P*_y_* are the cross-spectrum power density and *x*– and *y*-PSDEs, respectively.

Phase information between calcium and hemodynamics were investigated through the angle between the real and imaginary parts of the cross-spectrum power density., Phase averaging was performed in the complex plain to avoid issues with phase unwrapping. Owing to the features observed in the MSC plots (see Results), power was integrated over two frequency bands: between 0.5 Hz and 2 Hz and below 0.5 Hz.

### 4.15) Resting-state functional connectivity

Measures of resting-state functional connectivity (RSFC) were calculated using zero-lag correlation for all pixel pairs within the brain mask, as in previous reports [61, 79, 237, 238, 262]. Briefly, spontaneous activity was filtered to obtain frequencies over infraslow and delta frequency bands. Global signal regression was performed to reduce global coherence, resulting in increased spatial specificity of RSFC patterns [260, 263, 264]. Whole cortex correlation matrices were reordered according to the mouse brain atlas. Regional RSFC maps for frontal, cingulate, retrosplenial, and secondary motor (M2) were computed from the whole cortex by averaging all seed-based RSFC maps in each region (**Fig. 6**). Psychedelic-induced changes in RSFC were computed as RSFC post-injection minus RSFC pre-injection. Spatial similarity (Pearson correlation) and linear relationships (linear regression) between calcium– and hemodynamic reports of RSFC differences were computed (**Table 1**) as:

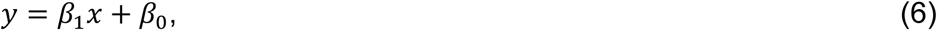

where *x* and *y* are vectorized RSFC difference matrices for calcium and HbT, respectively.

### 4.16) Inter-versus Intra-network functional connectivity alterations

Differential changes in inter– and intra-network connectivity is a common observation in studies examining the effects of psychedelics on systems-level brain function [16, 18, 80–82]. Modularity, which measures the strength of connections within versus between network communities, is often used to quantify these changes. In the context of psychedelics, it is often interpreted as changes in network integration [265]. However, while modularity is a commonly used measure to quantify these changes by assessing the strength of connections within versus between network communities, it is limited in topographical assessment. Furthermore, modularity obscures finer details of altered network organization, as it primarily focuses on community-level changes. In contrast, silhouette scores offer a more nuanced evaluation of network integration by measuring the similarity of individual nodes within clusters relative to their distance from other clusters.

To examine differences in RSFC structure before and after compound injection, RSFC matrices were computed for all pairwise comparisons within our field of view for each mouse before injection and averaged to yield a high-fidelity RSFC matrix (**Extended Data Fig. 10a**). Resting-state networks (RSNs) within this matrix were identified using Ward’s hierarchical clustering method [266, 267], which respects the intrinsic hierarchy within the data and has been previously applied to RSFC measures in humans [266, 268, 269]. Each branch of the resulting dendrograms (**Extended Data Fig. 10b**) represents a potential cluster. Moving from the bottom of the dendrogram to the top of the tree (corresponding to increased variance when merging), the threshold for defining a cluster becomes less stringent. Merging branches indicate that clusters have combined at less stringent criterion thresholds. A branch that remains separate over a longer vertical distance represents a more stable clusters across thresholds. A clustering threshold (black-dotted line, **Extended Data Fig. 10b**) was selected that aligns most closely with Allen-defined functional regions [270] and previous data-driven clustering results [248]. The resulting 13 clusters (**Extended Data Fig. 10c**) were used in subsequent analysis of RSFC changes through evaluation of silhouette scores.

Briefly, silhouette scores at each pixel quantifies how similar a pixel is to its own cluster compared to others (**Fig. E10d,i**), and like modularity [103], can be interpreted as a network measure related to cluster segregation. The algorithm computes a ratio between a measure of how similar a pixel’s RSFC map is to other pixel’s maps within the same cluster (cohesion) and how dissimilar the pixel’s map is from pixel’s maps in the nearest neighboring cluster (separation). Silhouette scores range from –1 to +1, such that +1 indicates a point is well-clustered (*i.e.*, integrated within its community and segregated from neighbors; high intra-connected, weak inter-connected), 0 indicates that the point is on or near the boundary between two clusters (equal intra– and inter-connectivity), and –1 indicates a point is misclassified and likely should be assigned to a different cluster than its own (weak intra-connected, strong inter-connected). Cohesion and separation were computed relative to each cluster’s centroid, computed as the average RSFC map within a cluster (vectorized as **C**_n_). Considering a pixel in the i-th pixel location (and vectorized RSFC Map, **r**_i_) and the k-th cluster, cohesion and separation are defined as:

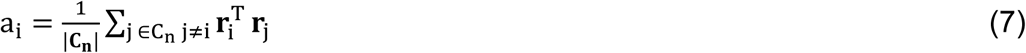

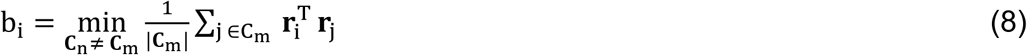

Then, silhouette scores (s_i_) were computed at each pixel:

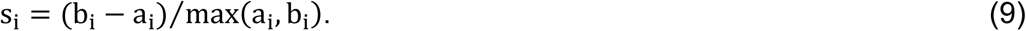

### 4.17) Statistics

Significant differences in movement and band-limited spectral content were determined using a Kruskal-Wallis one-way analysis of variance of the mean. For stimulus-evoked responses and associated HRFs and TFs, results are reported as medians and percentiles owing to the low number of stimulus presentations. Parameters extracted for evoked and resting-state activity and NVC were calculated as percent changes from pre-injection. Except where stated otherwise (owing to low n), significant effects of compound was determined using a Kruskal-Wallis one-way analysis of variance. Post-hoc significance was assessed using the paired Wilcoxon sign-rank test. Significant effects of time (*i.e.*, pre-versus post-injection comparisons) was assessed using the paired Wilcoxon signed-rank test. Multiple comparisons were corrected using the Bonferroni correction. When mean and standard deviation are used, effects of compounds were determined through one-way analysis of variance (ANOVA) and post-hoc t-tests. The effect of time (*i.e.*, pre-versus post-injection) was determined to be significant using paired t-tests and corrected for multiple corrections using Bonferroni correction. All tests were two-tailed.

For all RSFC measures, Pearson correlation coefficients were converted to Fisher z values prior to any mathematical operation and converted back to Pearson correlations for visualization. Statistical differences in seed-based RSFC were evaluated on a cluster-wise basis [271, 272], with a clustering threshold of 12 contiguous pixels.

## Supporting information

All supplemental figures and text

## Acknowledgements

This work was supported by National Institutes of Health grants R01NS126326 (A.Q.B.), R01NS102870 (A.Q.B.), RF1AG07950301 (A.Q.B.), R01NS117899 (J.G.M.), R01NS135401 (J.G.M.), T32DA007261 (J.S.S), R25 MH112473, the Taylor Family Institute for Innovative Psychiatric Research, the Center for Brain Research in Mood Disorders (CBRiMD) and the Center for Holistic Interdisciplinary Research in Psychedelics (CHIRP) at Washington University School of Medicine in St. Louis (J.S.S., G.E.N), F99NS139512 (J.A.P.C.), T32EB014855 (J.A.P.C.), T32NS121881 (O.J.K.). We thank Ryan Raut and Timothy Laumann (Washington University in St. Louis) for helpful discussions and advice. We would also like to express our sincere gratitude to the mice for their vital contributions to this study, which were essential for the findings presented in this paper. We would like to acknowledge the Osage Nation, Missouria, Illinois Confederacy and many other tribes as the ancestral, traditional, and contemporary custodians of the land where this work was conducted.

## Author contributions

Experimental Design: J.A.P.C., J.S.S, J.G.M., and A.Q.B.

Technical Recourses: J.A.P.C., O.J.K, C-.C.K., X.W., J.G.M., and A.Q.B.

Animal Preparation and Surgeries: J.A.P.C., O.J.K, C-.C.K., A.R.B

Data Acquisition: J.A.P.C., O.J.K, C-.C.K.

Data Analysis: J.A.P.C., O.J.K, C-.C.K., J.S.S

Project Supervision: J.S.S, J.G.M., and A.Q.B.

Writing—Original Draft: J.A.P.C, A.Z.S, J.S.S, J.G.M., and A.Q.B.

Writing—Review & Approval: J.A.P.C, O.J.K, C-.C.K., X.W., A.R.B., G.E.N., A.Z.S, J.S.S, J.G.M., and A.Q.B.

Financial Support: J.A.P.C., G.E.N., J.S.S., J.G.M., and A.Q.B.

## Figures and Tables

**Extended Data Figure 1).**
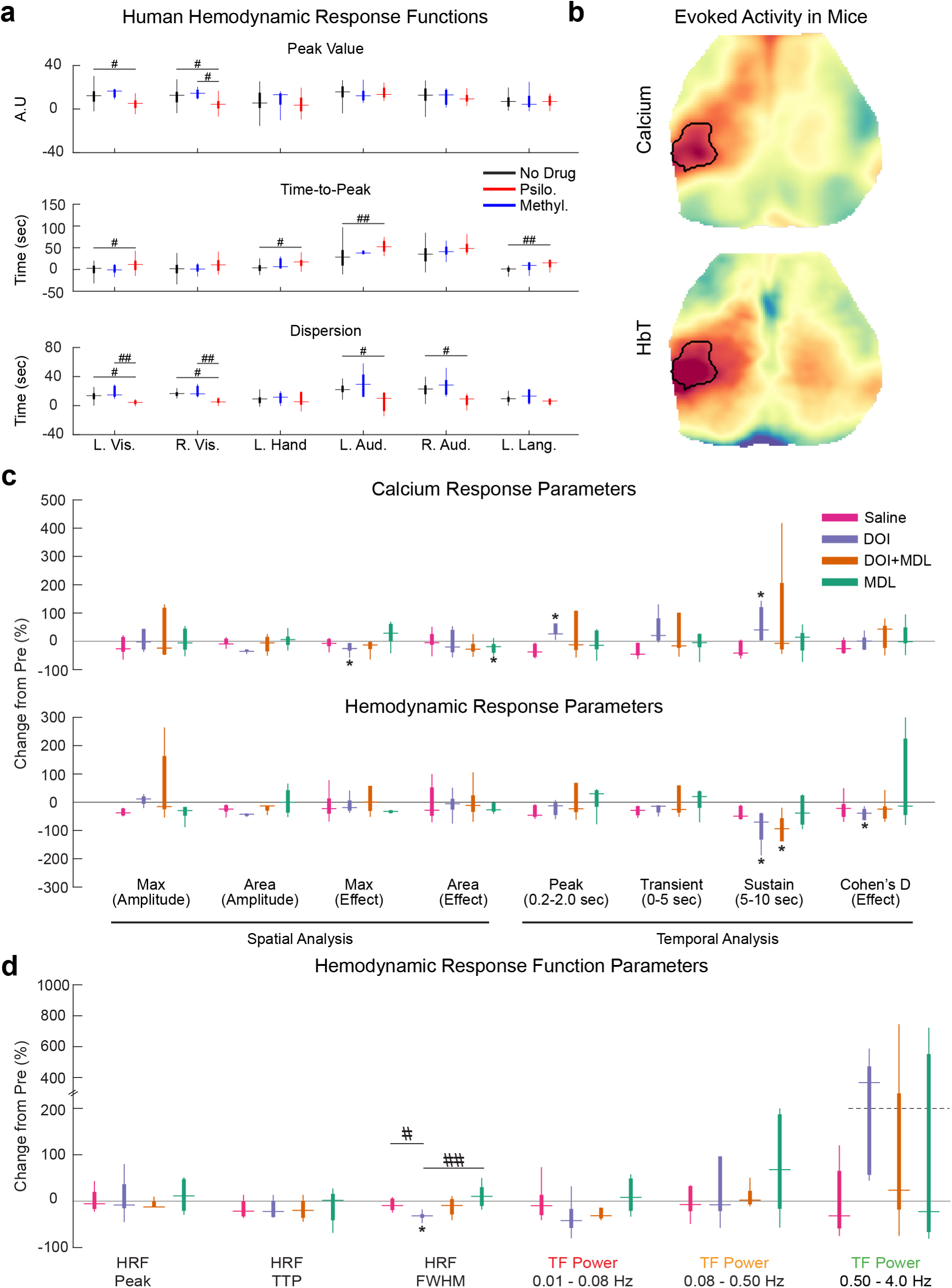
Quantification of task-based fMRI and WFOI changes induced by hallucinogenic 5-HT2A receptor agonism. a) Quantified changes in hemodynamic response functions in humans. Generalized linear models were fit for task-evoked activity in left/right visual cortex, left hand, left/right auditory cortex, and left Wernicke’s area (left language). HRFs were fit using double gamma basis functions characterized by three-parameters: Peak Value (P), Dispersion (D), and Time to Peak (TTP). Statistical differences in each parameter across conditions were assessed *via* 2-way ANOVA; significant effects were evaluated post-hoc *via* t-tests. p-values are indicated such that #<0.05, ##<0.01, ###<0.001. **Top panel:** Peak value. Peak value decreased in in left and right visual regions under psilocybin conditions (ANOVA, p=0.03 and p=0.020, respectively). **Middle panel:** Time-to-peak. Time-to-peak decreased in all regions except the right visual region (ANOVA, left visual, p=0.03 left hand, p=0.006; left auditory, p=0.002; right auditory, p=0.044; left language, p=0.007). **Bottom panel:** Dispersion decreased in the left and right visual regions, and left and right auditory regions (ANOVA, p=0.007, p=0.004, p=0.021, p=0.022, respectively). **b.) Pre-injection stimulus.** Pre-injection effect maps for all saline, DOI (high dose), DOI (high dose) + MDL and MDL (n=32; 8 mice ·4 compound conditions) were combined and averaged to create a single, high-signal-to-noise ratio response effect map. To define a consistent response criterion across all mice and compounds, effect maps were thresholded at 75% of the maximum value (top 25%). This created a shared region-of-interest (ROI) for subsequent analysis across all mice. **c) Quantification of calcium and hemodynamic response dynamics.** Within the ROI, response amplitude, area (*i.e.*, number of pixels over 75% of the maximum), and Cohen D’s effect size were tabulated and visualized. Temporal dynamics were parameterized by extracting the peak value over the inter-stimulus period (0.2-2.0 seconds for calcium and 2.2-4.2 seconds for HbT), the area under the curve during the transient response (0-5 seconds) and the sustained response (5-10 seconds), and Cohen’s D effect. **d) Parameter quantification of stimulus evoked neurovascular coupling.** For stimulus-based hemodynamic response functions (HRFs), the peak value (peak), time to peak (TTP), and full-width-at-half maximum (FWHM) were computed. After injection of DOI, FWHM decreased (–31.4% [–36.0%, – 27.8%]; pre-versus post-injection: p=0.031). For transfer functions (TF), band-limited frequency transduction was parameterized in three frequency bands: infraslow activity (ISA, 0.01 Hz –0.08 Hz), intermediate band (0.08-0.50 Hz), and delta-band (0.5 Hz-4.0 Hz). All outliers (>99.3^rd^ percentile) were excluded for visualization purposes only. Changes from pre-injection to post-injection was assessed through Wilcoxon’s sign-rank tests. *<0.05, **<0.01, ***<0.001. Differences (pre-minus post-) across drugs of compound were tested using a one-way Kruskal-Wallis and post-hoc Wilcoxon’s sign-rank tests. #<0.05, ##<0.01, ###<0.001. Multiple comparisons were corrected for using the Bonferroni method.

**Extended Data Figure 2.**
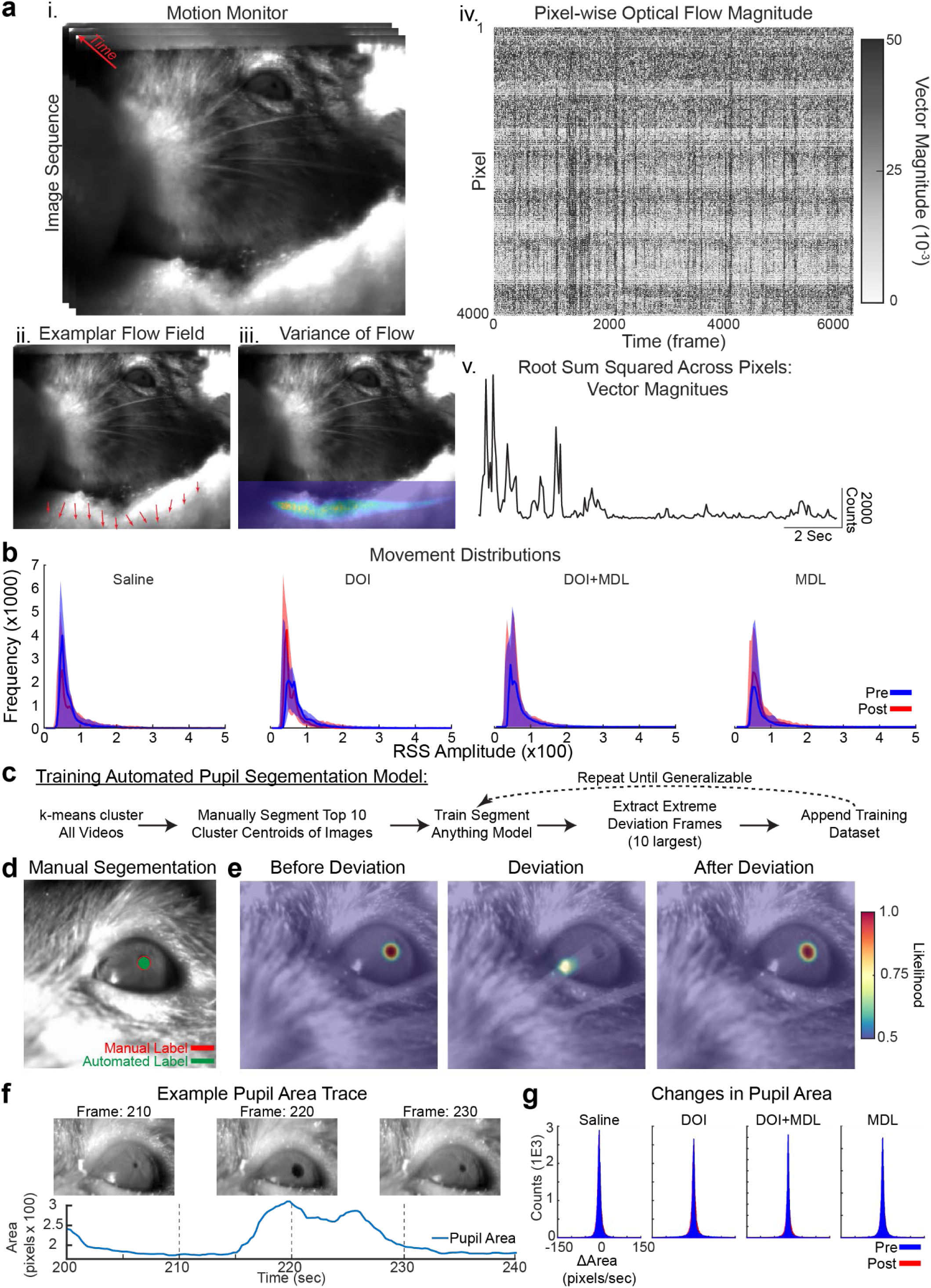
DOI does not alter motion or pupil dynamics. Two CMOS cameras (frame rate: 20 Hz, movement field-of-view ∼70mm-by-70mm, pupil field-of-view ∼8mm-by-8mm), were placed in front of the mouse to monitor body movement and pupil dynamics. Illumination was delivered through a 750 nm, near infrared (NIR), LED, and NIR polarizers were used to minimize specular reflection. Image sequences of mouse movement and pupil dynamics were time locked to WFOI. **a**) **Motion monitoring using optical flow.** Movies of the front of the mouse were recorded (**i**) and optical flow (OF) estimates were generated using the Lucas-Kanade method with a 1-second temporal window for all imaging sequences (**ii**); OF resulted in spatiotemporal series of vectors for which each pixel and frame represents the spatiotemporal gradient. The temporal variance of vector magnitudes across time demonstrates OF sensitivity to motion of the felt supporting the mouse (**iii**). For each pixel, the vector magnitude can be computed, with higher values corresponding to larger movement (**iv**). The root-sum square across all frames was computed to yield a single time trace of estimated movement (**v**). **b) Distributions of OF-derived movement measures**. No differences were found between compounds (as assessed by differences between post-minus pre-injection; Kruskal-Wallis: p=0.262). **c**) **Simplified schematic of the automated pupil segmentation algorithm. d**) **Example of a user-annotated pupil contour and the automated pupil contour using our algorithm.** As demonstrated, automated segmentation often surpassed manually segmented precision. **e**) **Example of the automated outlier detection algorithm using the center-of-mass of the likelihood distribution.** Large deviations in likelihood distributions are attributed to artifact due to the slow rate of pupil diameter fluctuations compared to the imaging frame rate (20 Hz). Outlier frames, along with the frames immediately before and after, were labeled and appended to the original training dataset to increase robustness of the algorithm. **f**) **Example frames and time course of pupil area changes during imaging.** Large changes in pupil diameter are reflected in changes in pupil area. **g**) **Distributions of pre– and post-injection changes in pupil area for each compound condition.** No differences in pupil dynamics before and after compound injection were observed for any condition.

**Extended Data Figure 3).**
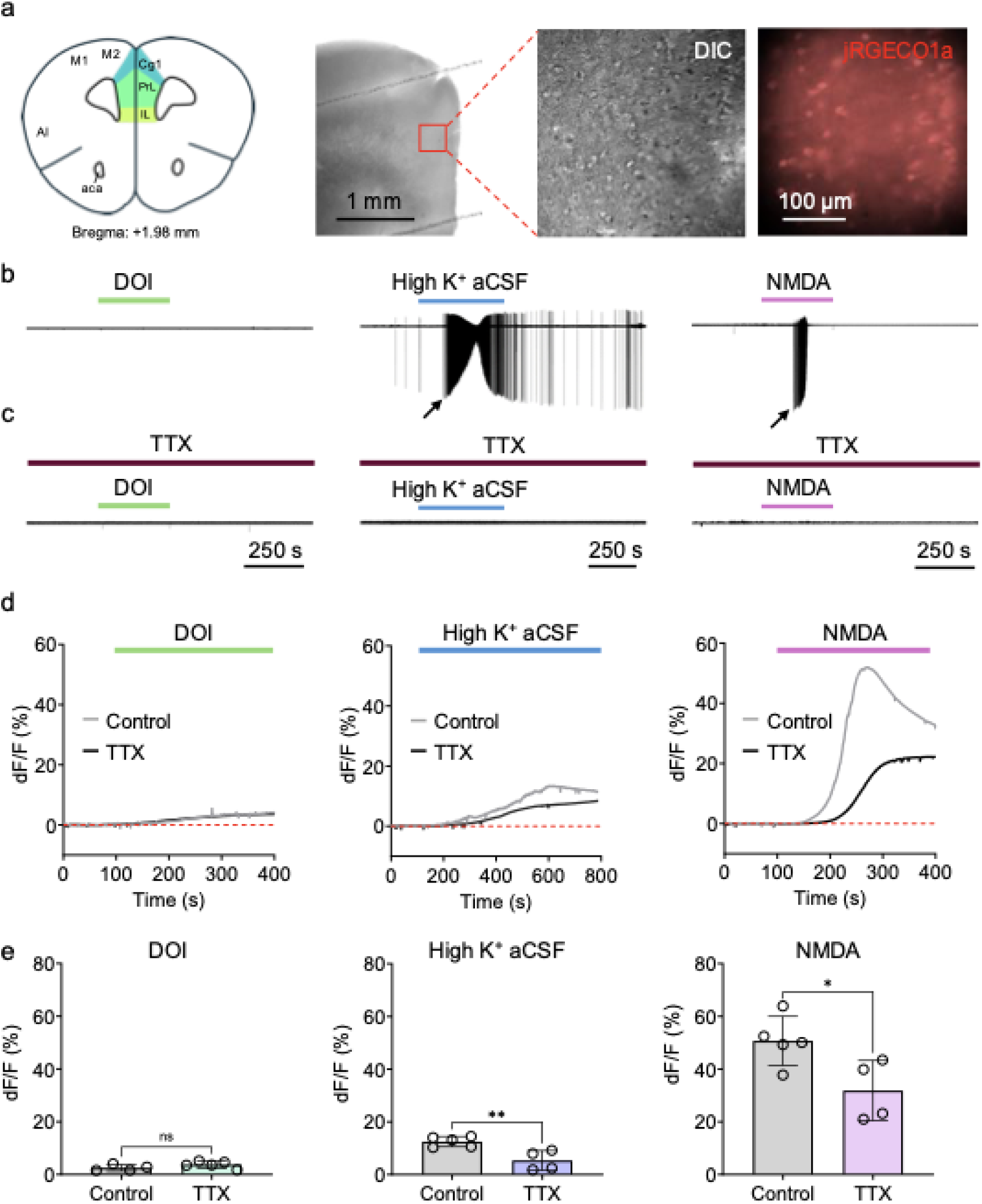
Limited contribution of DOI to action potential-independent calcium fluctuations. *Ex vivo* recordings of global calcium signal in acute brain slice upon DOI (10 μM), high K^+^ aCSF (15 mM) or NMDA (50 μM) application. **a) Experimental arrangement.** *Ex vivo* recordings were conducted on pyramidal cells across the prefrontal cortical region (*e.g.*, cingulate, prelimbic and infralimbic cortices). **b) Direct activation of 5-HT_2A_ receptors does not drive action potentials.** Representative cell-attached recordings demonstrate low spontaneous activity of cortical pyramidal cells in acute brain slice preparation. High K^+^ aCSF and NMDA, but not DOI, result in saturated neuronal firing (black arrows). **c) TTX fully suppresses evoked action potentials.** These bursts were completely suppressed by tetrodotoxin (TTX; 1 µM) application. (Control: DOI: 6 cells, High K^+^ aCSF: 8 cells, NMDA: 5 cells. TTX: DOI: 4 cells, High K^+^ aCSF: 4 cells, NMDA: 4 cells.) **d) Small, but steady TTX-resistant global calcium increase upon DOI application.** Representative traces show robust, 755 long-lasting enhancement of global calcium signal by High K+ aCSF and NMDA compared to DOI**. e) TTX does not alter DOI-induced calcium increase.** TTX pretreatment significantly suppressed the High K^+^– and NMDA-induced increase in calcium signal but did not alter the subtle DOI-mediated calcium enhancement. (Student’s t-test, DOI: 2.50 ± 1.18 vs. 3.73 ± 1.30%, p=0.1878; High K^+^ aCSF: 12.50 ± 1.95 vs. 5.34 ± 3.78%, p=0.0075; NMDA: 50.23 ± 10.70 vs. 31.85 ± 11.44%, p=0.0293). Abbreviations: aca, anterior commissure (anterior part); AI, anterior insular cortex; Cg1, cingulate cortex (area 1); IL, infralimbic cortex; M1, primary motor cortex; M2, secondary motor cortex; PrL, prelimbic cortex.

**Extended Data Figure 4).**
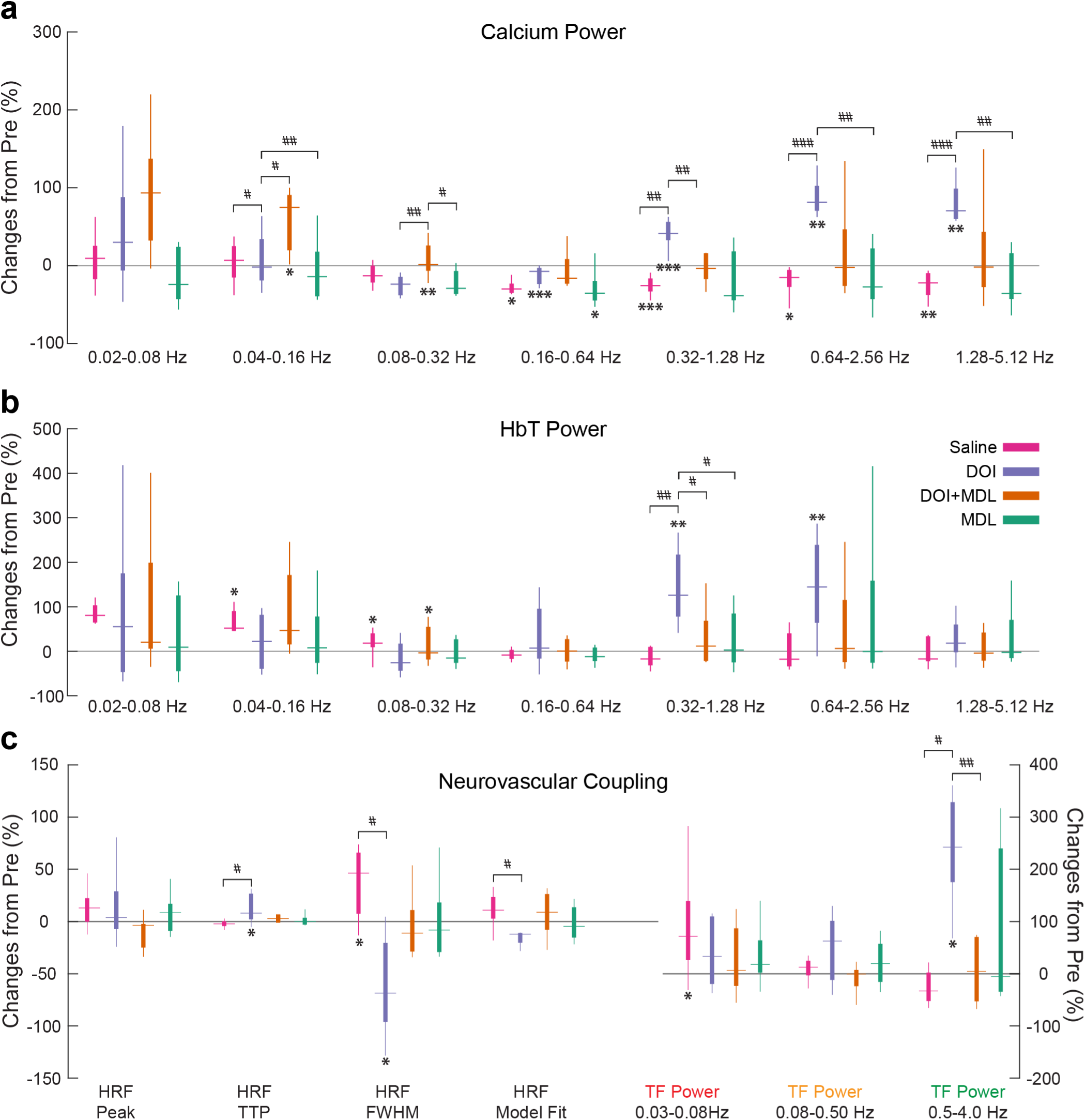
Hallucinogenic 5-HT_2A_ receptor agonism changes resting-state power and neurovascular coupling. **a-b**) Power spectral density estimates were integrated over double-octave bins and compared between compounds for both calcium (**a**) and hemodynamic (**b**) activity. The lowest frequency bin was limited by the lower limit of 0.02 Hz due to the window length used to segment resting-state from stimulus-evoked data. All measures were averaged over the cortex and comparisons were made between pre– and post-injection and across compounds (post-minus pre-) using t-tests (*<0.05, **<0.01, and ***<0.001) and one-way ANOVAs with t-test post-hoc tests (#<0.05, ##<0.01, ###<0.001), respectively. Multiple comparisons were corrected for using the Bonferroni method. **c.) Neurovascular coupling quantification.** Regional hemodynamic response functions (HRFs) were characterized by their peak value (peak), time to peak amplitude (TTP), and full width at half maximum (FWHM). These parameters were averaged across regions and compared. Predicted hemodynamic activity at each region was calculated by convolving each region’s HRF with the measured calcium activity in that region. Predicted and measured hemodynamic activity were compared *via* Pearson r correlation coefficient as a measure of model fit. Band-limited frequency transduction was calculated over three frequency bands: infraslow activity band (ISA, 0.03-0.08Hz), intermediate activity band (0.08 Hz-0.50 Hz), and delta activity band (0.5 Hz-4.0 Hz). The lowest frequency bin of the TF (*i.e.*, 0.03Hz) was determined by 30 second window length used for estimating HRFs.

**Extended Data Figure 5).**
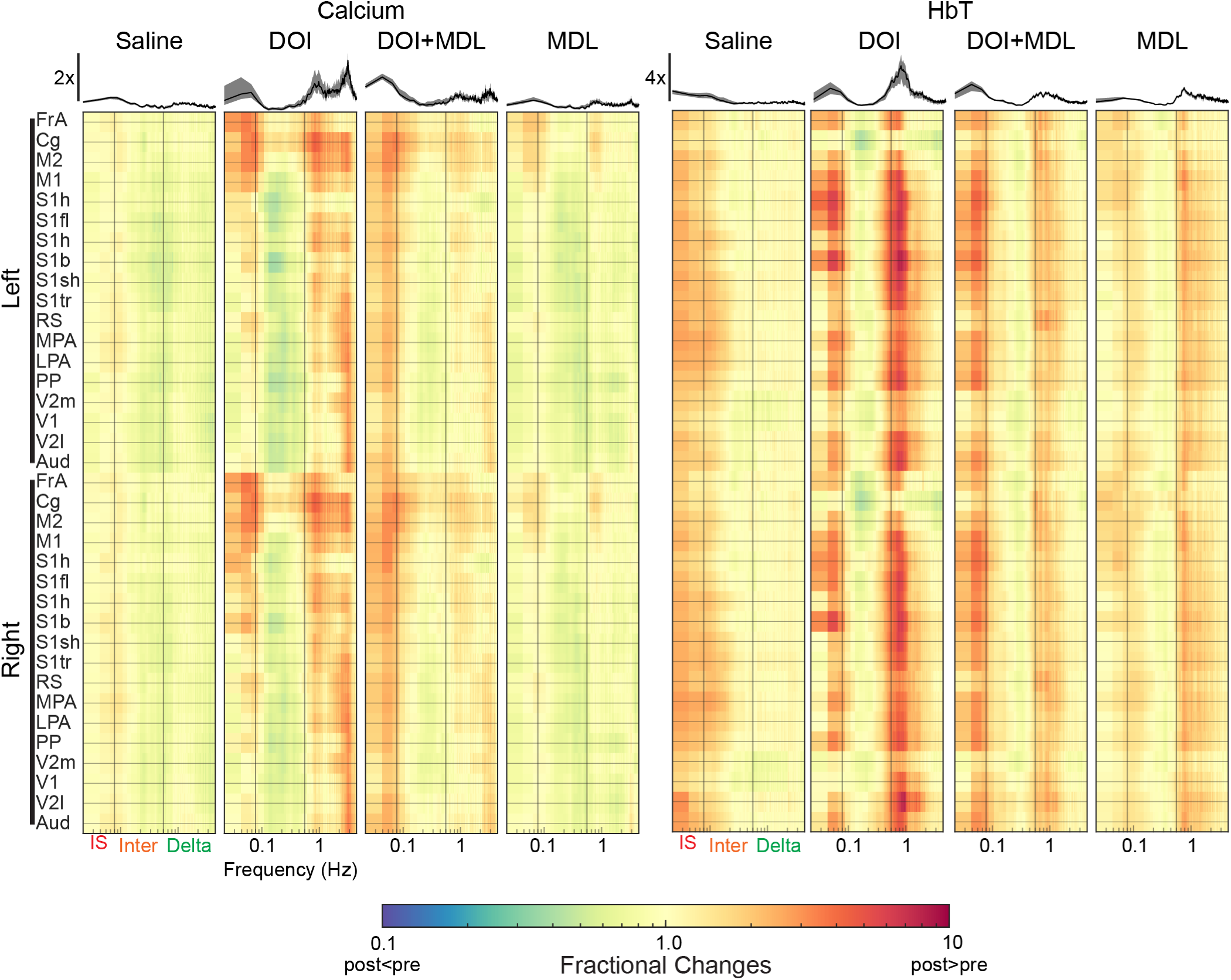
Fractional changes in broad-band activity attributed to hallucinogenic 5-HT_2A_ receptor agonism. Regional power spectral density estimates (see Fig. 4c) were characterized over the following frequency ranges: infraslow (ISA): 0.01 Hz-0.08 Hz; intermediate: 0.08 Hz-0.50 Hz; and Delta: 0.5 Hz-4.0 Hz (Fig. 4c). Fractional changes in regional power spectra are displayed as the ratio of post-injection values divided by pre-injection values. DOI differentially affected the spectral content of calcium (**right**) and hemodynamic (**left**) activity in a region-dependent manner. *<u>ISA</u>*: DOI increased calcium ISA power in frontal, cingulate, and motor cortices while hemodynamic ISA activity increased primarily in somatosensory regions. *<u>Intermediate</u>*: Calcium activity decreased in all cortical regions excluding frontal and cingulate. In contrast, hemodynamic activity decreased in frontal and cingulate cortex. *<u>Delta</u>*: Hemodynamic activity (right panel) exhibited increased delta-band activity in nearly every cortical region examined expect for in cingulate and retrosplenial cortex. Increases in faster signaling were also present in calcium signals (left panel). Across all frequencies examined, MDL largely reversed the effects of DOI. Saline and MDL minimally altered calcium and hemodynamic spectral content. Averaged fractional changes over the cortex are visualized on top of each column and plotted as mean +/-standard deviation across mice.

**Extended Data Figure 6).**
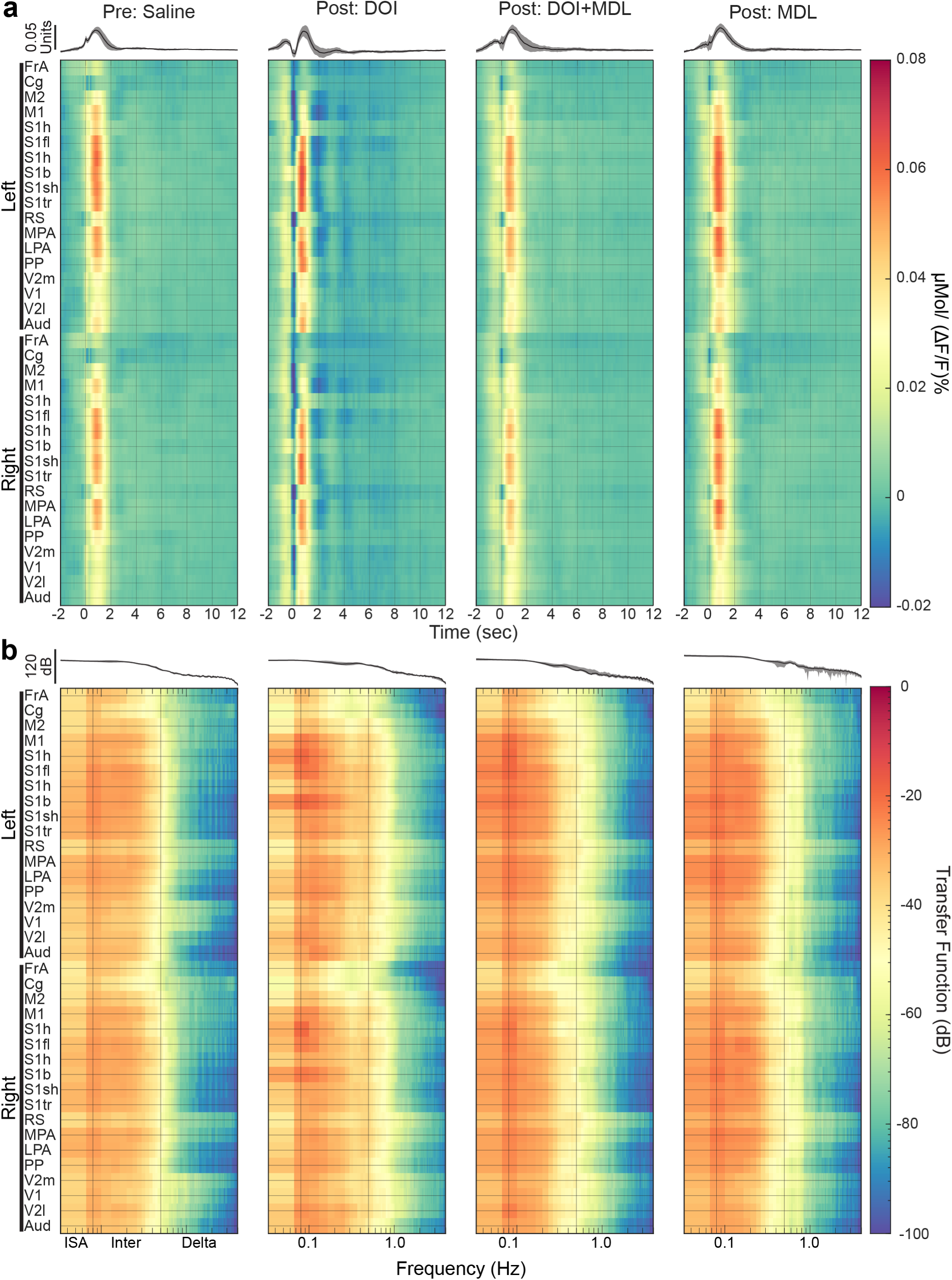
Hallucinogenic 5-HT2A receptor agonism alters region-specific NVC. a) Regional HRFs. Estimated HRFs for each region were averaged across mice for each compound. Regions are organized from anterior-to-posterior and from left-to-right. Prior to injection of any compound, NVC differed (parameterized in Fig. 4c) across the cortex. For instance, somatosensory and parietal regions exhibited the highest degree of coupling (*e.g.*, peak value), while frontal and cingulate regions exhibited more modest coupling. DOI caused a zero-lag notch across the cortex (*i.e.*, the acausal feature seen in Fig. 4a, black arrow), strongest in secondary motor, retrosplenial, and visual regions. Additionally, DOI increased coupling in the somatosensory and parietal regions. These effects were dampened when DOI was administered with MDL. Averaged HRFs over the cortex are visualized on top of each column and plotted as mean +/-standard deviation across mice. The color axis has units of μMol/(ΔF/F%). **b) Regional TFs.** Prior to compound administration, frequency dependent calcium-to-hemodynamic transduction differed across the cortex. For instance, cingulate and retrosplenial regions exhibited larger broad-band transduction compared to other brain regions. DOI was associated with broader band (power above 0.5 Hz, with a ∼1 Hz peak) transduction across the cortex. These effects were largely reversed when DOI was co-administered with MDL. Averaged TFs over the cortex are visualized on top of each column and plotted as mean +/-standard deviation across mice. TFs have units of (μMol/(ΔF/F%))^2^/Hz).

**Extended Data Figure 7).**
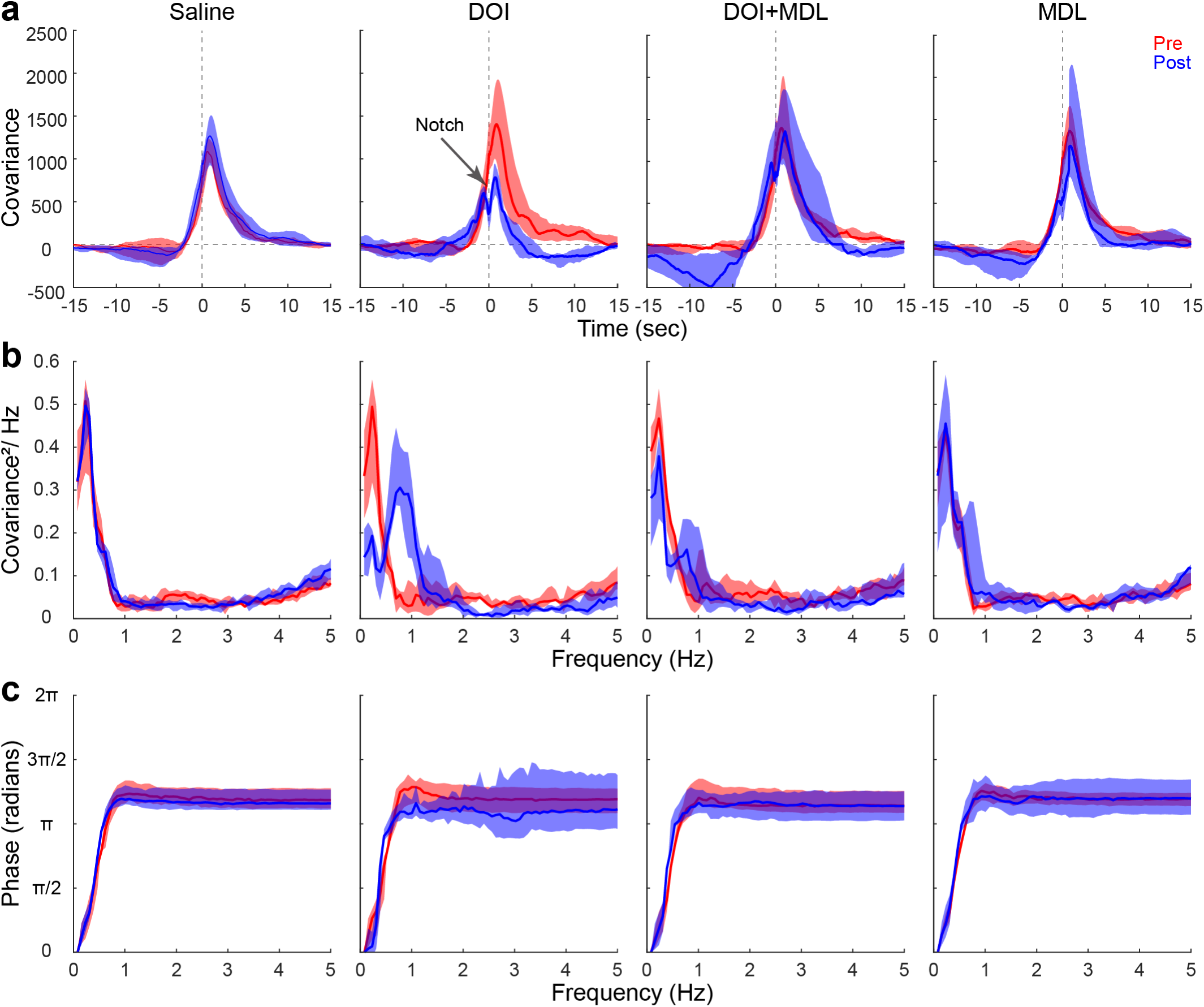
Lagged cross-covariance and coherence between calcium and total hemoglobin recapitulate DOI-induced changes in neurovascular coupling. a) Cross-covariance between calcium and total hemoglobin. The lagged cross-covariance function (CCF) between unfiltered, resting-state calcium and hemodynamic activity was computed for each cortical region over non-overlapping, 30 second windows (*n* = 54, as performed with HRFs) for each mouse. Pre-injection CCFs display a simple, quasi-causal (*i.e.*, calcium predominantly leading hemodynamics) relation. DOI significantly decreased synchrony (decreased peak value, –51% [–66%, –13%] pre-versus post-injection: p=0.039; Kruskal-Wallis: p=0.004; saline versus DOI: p=0.020; DOI versus DOI+MDL: p=0.004; DOI versus MDL: p<0.001). As observed in global HRFs (Fig. 3a), the DOI included a zero-lag notch in the CCF (black arrow; see main text, **Section 4.3.1**). Covariance has units of μMol(ΔF/F%). **b) Coherence between calcium and hemodynamics.** To better understand the frequency content contained in the cross-covariance functions, magnitude squared coherence (MSC) between calcium dynamics and hemodynamics were computed for each region and time window. Before injection of compounds, MSC exhibited a peak at 0.2 Hz, demonstrating the band-limited nature of NVC. After injection of DOI, a coherence peak ∼0.8 Hz (∼0.5 Hz half-bandwidth) emerged (0.5 Hz – 2.0 Hz: +170% [110% 230%]; pre-versus post-injection: p=0.008; Kruskal-Wallis: p=0.004; saline versus DOI: p=0.009; DOI versus DOI+MDL: p=0.030; DOI versus MDL: p=0.025) and the 0.2 Hz peak largely diminished (<0.5 Hz: –47% [–53%, 49%], pre-versus post-injection: p=0.008). This phenomenon signifies a DOI-induced shift in the coherent frequencies contained in both neuronal and hemodynamic activity. As demonstrated in other analytical measures, DOI+MDL largely reversed the effects of DOI alone. **c) Phase relation between calcium and hemodynamics.** For all compounds, phase exhibited a calcium lag-lead relation, reflected in the positive slope at frequencies below ∼1 Hz. All data are presented mean +/-standard deviation across mice.

**Extended Data Figure 8).**
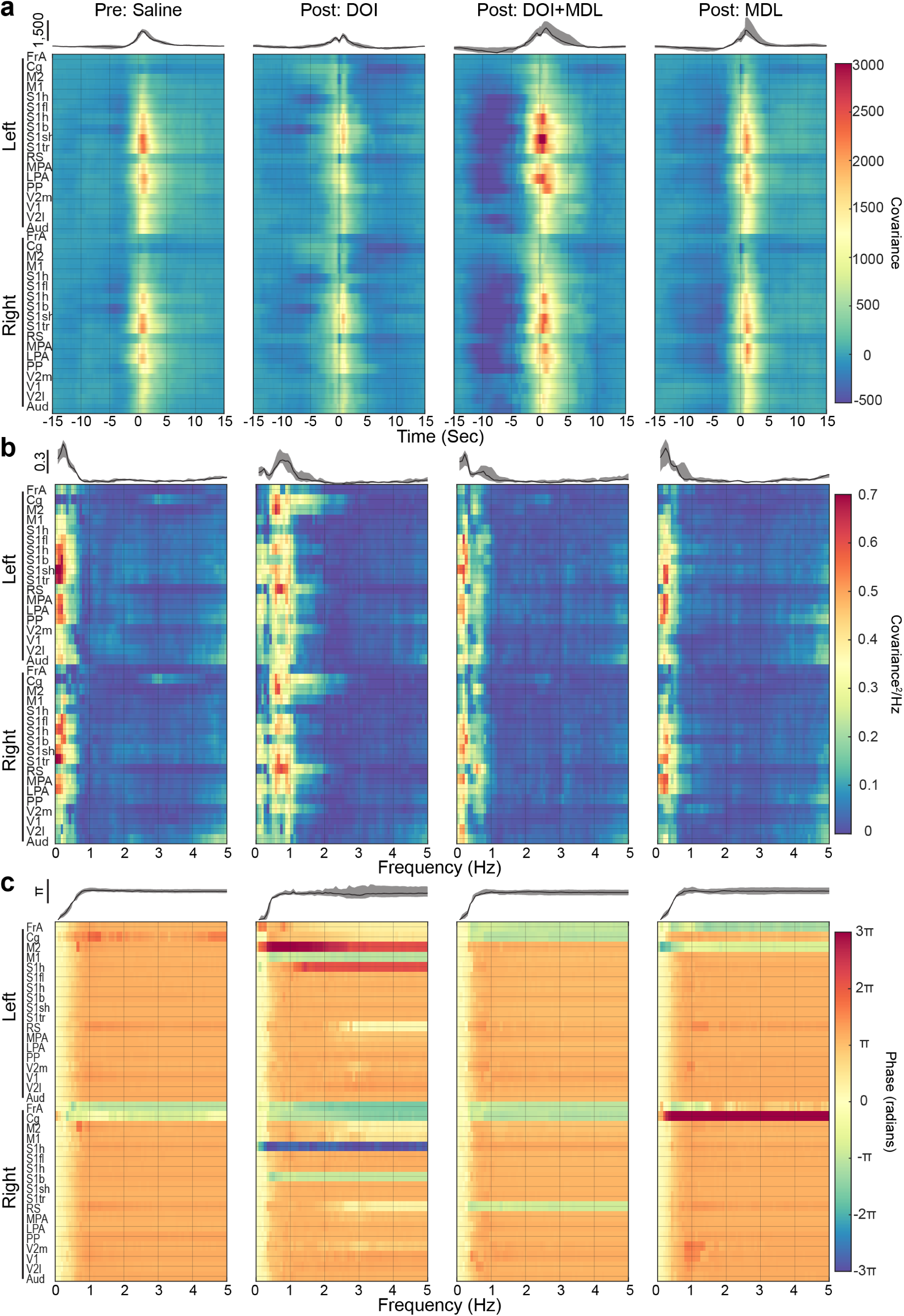
Regional lagged cross-covariance and coherence. a) Lagged cross-covariance between calcium and hemodynamics reveal regional-specific contribution to global cross analysis. The lagged cross-covariance function (CCF) between resting-state calcium and hemodynamic activity in each region was calculated as in **Fig. E7**. Before injection of compound (left), coupling (*i.e.*, peak value) was strongest in somatosensory and parietal regions and weakest in frontal, cingulate, retrosplenial, and motor regions. As evident in regional HRFs (**Fig. E6a**), DOI decreased cortical-wide coupling strength and caused a sharp notch at zero-lag ubiquity across the cortex. For example, the notches were most evident in secondary motor, retrosplenial, and visual regions. Post-injection DOI CCFs also decayed faster in all cortical regions, with stronger decreases in the cingulate and retrosplenial regions. DOI+MDL predominantly reflected pre-injection CCFs, except for decreases in somatosensory and parietal regions at negative lag times (time<0 seconds). Averaged CCFs over the cortex are visualized on top of each column and plotted as mean +/-standard deviation across mice. Covariance has units of μMol·(ΔF/F%). **b) Coherence between calcium and hemodynamics.** Magnitude-squared coherence (MSC) was calculated in each region. As observed in pre-injection global measures, there was strong coherence in lower frequencies (<0.5 Hz), which was largest in somatosensory and parietal regions and weakest in frontal, cingulate, motor, and retrosplenial regions. DOI decreased low-frequency coherence in a region-specific manner, with the largest reductions in higher order regions (*e.g.*, cingulate and retrosplenial). An emergent peak in MSC at ∼0.8 Hz (**Fig. E7b**) was primarily in higher order regions such as cingulate, secondary motor, and retrosplenial cortices. Notably, before injection of DOI, these regions showed low, broad-band coherence. Regional coherence under DOI+MDL reflected pre-injection characteristics. Averaged MSC over the cortex are visualized on top of each column and plotted as mean +/-standard deviation across mice. **c) Phase relation between calcium and hemodynamics.** Before injection, phase relations did not show regional variation (left). After injection of DOI and DOI+MDL, most regions matched their pre-injection characteristics, except for frontal, cingulate, and retrosplenial regions, where phase slopes reversed (*i.e.*, yellow pre-injection, greens and blue post injection), revealing that hemodynamics lead calcium in these regions (*i.e.*, vasculo-neural coupling). MDL did not fully reverse this phenomenon in all regions.

**Extended Data Figure 9).**
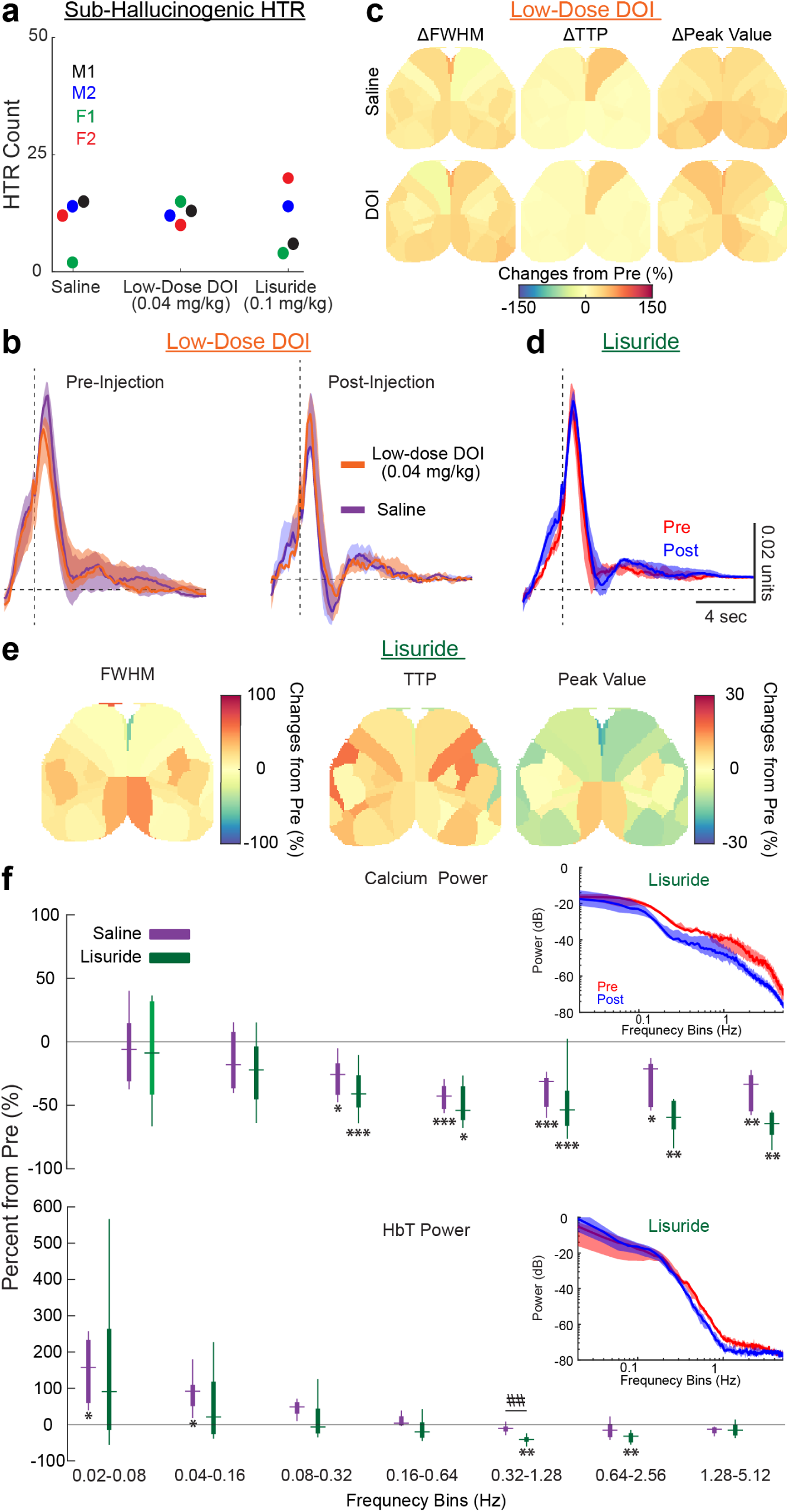
Non-hallucinogenic doses of DOI and the non-hallucinogenic 5-HT_2A_R agonist, Lisuride do not alter NVC. **a**) HTR results from sub-hallucinogenic (low dose) DOI (0.04 mg/kg) and the non-hallucinogenic, psychedelic ligand, Lisuride (0.1 mg/kg). No differences in HTRs were observed between groups (KW>0.99). HTRs were recorded for 30 minutes. **a-f) WFOI imaging of non-hallucinogenic ligands.** Two groups of Thy1-jRGECO1a mice were imaged under resting-state conditions for 30 minutes before and 30 minutes after injections. Cohort 1 (4 males, 5 females) received saline and Lisuride (0.1 mg/kg); cohort 2 (2 males, 2 females) received saline and a sub-hallucinogenic dose of DOI (low-dose DOI, 0.04 mg/kg. **a**) Low-dose DOI did not alter NVC **b) Topographical changes in HRF parameters** full-width-at-half-maximum (FWHM), time-to-peak (TTP), and peak value are visualized. Sub-hallucinogenic doses of DOI also did not significantly alter any regional HRF parameters compared to Saline. **c) Global power spectral density estimates before and after injection of Lisuride** and **d) integrated, band-limited power over double octave frequency bins**. Band-limited power is consistently decreased after injection of both saline and Lisuride, like results reported in **Fig. E5a-b**. Lisuride modestly affected power over the 0.32-1.28 Hz bin (p=0.009). **e) Hemodynamic response functions before and after Lisuride injection.** No significant changes to hemodynamic response functions were observed after the injection of Lisuride. **f) Regional changes NVC following low-dose DOI.** Parameterized hemodynamic response functions reveal no differences in regional NVC after injection of low-dose DOI.

**Extended Data Figure 10).**
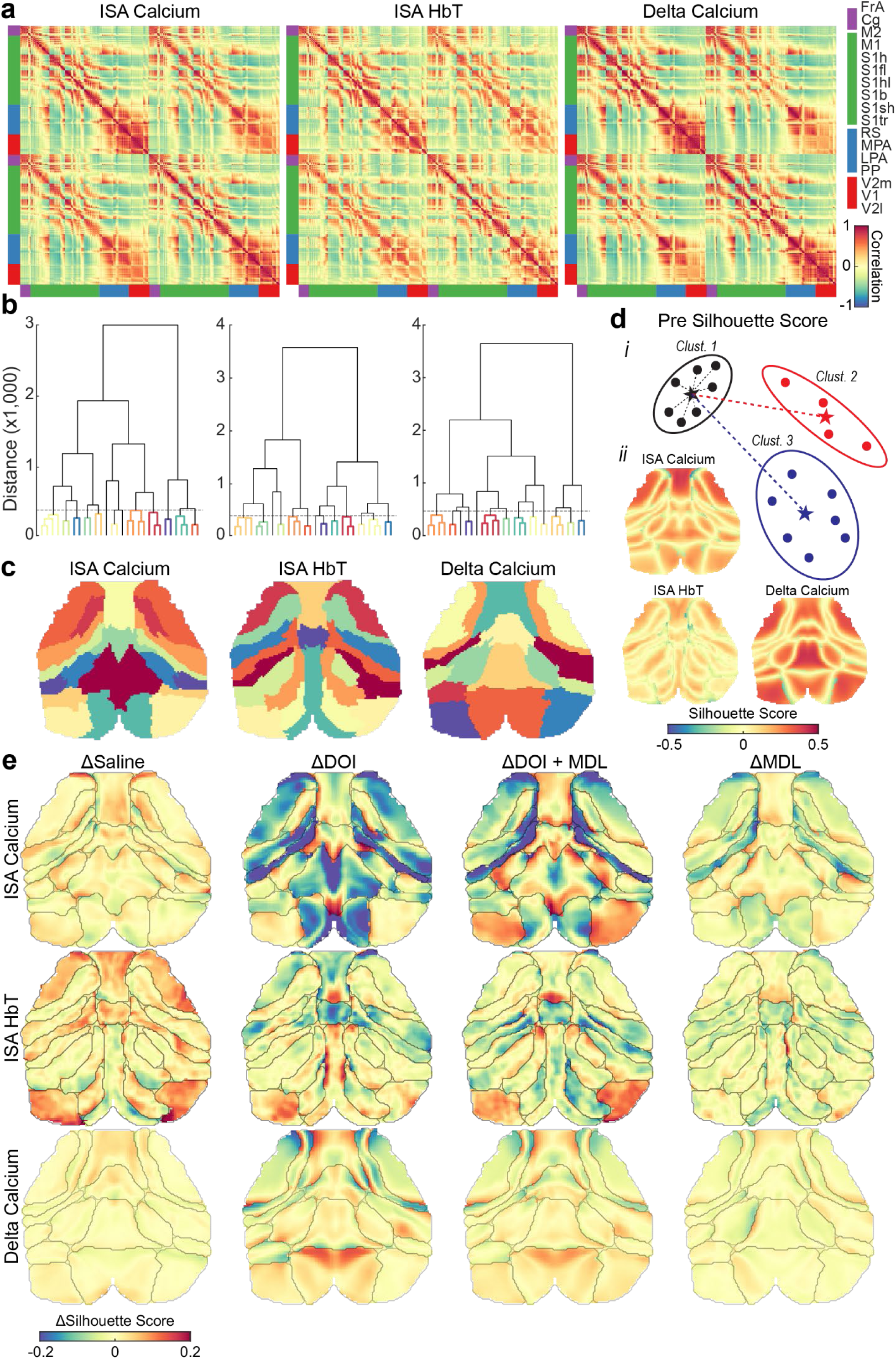
Hallucinogenic 5-HT_2A_ receptor agonism alters inter-versus intra-network promiscuity. **a**) **Whole-cortex RSFC matrices.** Average, pre-injection, whole-cortex RSFC matrices across mice for infralow (ISA; 0.01-0.08 Hz) calcium and HbT activity and delta (0.5Hz-4Hz) calcium activity. Matrices represent all pixel-pair comparisons and are organized by network assignment. **b) Hierarchical clustering of RSFC.** RSFC matrices were hierarchically clustered using Ward’s method and the hierarchy was visualized *via* dendrograms. **c**) **Resultant clusters at specific hierarchical level.** A distance threshold was selected to create a parcellation containing 13 parcels. These parcels were used to assess inter– and intra-regional relationships. **d**) **Pre-injection Silhouette scores.** Silhouette scores were computed at each pixel to assess the ratio between cohesion (intra-cluster distance metric) and separation (inter-cluster metric). Silhouette scores range from –1 to +1, such that +1 indicates a point is well-clustered and far from neighboring clusters (highly intra-connected, weakly inter-connected), 0 indicates that the point is on or near the boundary between two clusters (equally intra– and inter-connected), and –1 indicates a point is misclassified and likely should be assigned to a different cluster than its own (weakly intra-connected, strongly inter-connected). **i**) A graphical demonstration of cohesion (black cluster) and separation to the nearest cluster (red cluster). **ii**) Pre-injection silhouette scores for each species and frequency band. Importantly, we note that boundaries have scores of zero, reflecting the nature of boundaries (*i.e.*, equally likely to belong to neighboring clusters). **e**) **Changes in Silhouette scores after compound injection.** DOI predominantly decreases ISA calcium scores everywhere (regions are less intra-connected and more inter-connected), an effect not fully reversed in under DOI+MDL. Importantly, scores in the cingulate and retrosplenial regions (constellations of the mouse default mode network) differentially changed, with the cingulate marginally increasing and the retrosplenial largely decreasing. This effect is drastically decreased in ISA HbT-based scores. Scores for delta calcium activity under DOI reveal that DOI primarily alters network boundaries more than affecting inter-or intra-network interactions.

