## Supplementary material for "Psychedelic 5-HT_2A_ receptor agonism: neuronal signatures and altered neurovascular coupling": All supplemental figures and text

### Supplemental Results

*N1) Validation of NVC estimates*

Lagged cross-covariance and coherence were calculated to validate observed changes in resting-state NVC between calcium and total hemoglobin.

*N1.1) Global lagged cross-covariance analysis*

Cortical averaged cross correlation functions (CCFs) were compared before and after injection compounds (**Extended Data Fig. 7a**). Before injection (pre-injection), global CCFs reflected a simple, causal (*i.e.*, calcium leading hemodynamics) relation. Injection of saline and DOI+MDL (post-injection) minimally altered this relationship. As observed in the HRFs of **Figs. 5** and **Extended Data Fig. 6** DOI decreased positive lag lagged synchrony (*i.e.*, correlation), as reflected in a decreased CCF value. Moreover, the global CCF decayed faster after DOI. Importantly, as observed in HRFS, the DOI CCF exhibited a notch at zero lag and a resultant “acausal” (occurring before time zero) peak (**Extended Data Fig. 7a**, middle, black arrow).

Magnitude squared coherence functions (MSC) were calculated to assess the frequency content present in global CCFs (**Extended Data Fig. 7b**). Before injection of compounds, the global MSC exhibited a peak at 0.2 Hz, indicating the band-limited nature of NVC. Following the injection of DOI, a peak in coherence emerged at approximately 0.8 Hz (with a half-bandwidth of ~0.5 Hz), while the 0.2 Hz peak significantly diminished, as indicated by the decreased amplitude of slow oscillations in the time-domain results (*i.e.*, CCFs, **Extended Data Fig. 7a**). This phenomenon signifies a DOI-induced shift in the coherent frequencies contained in both neuronal and hemodynamic activity. These aspects of NVC alterations from DOI alone were not present when DOI was co-administered with MDL.

To quantify the above observations, coherence was integrated from 0.5 Hz to 2.0 Hz or below 0.5 Hz for all compounds. Saline did not alter coherence between calcium and hemodynamic activity. There was a decrease in low frequency (<0.5 Hz) coherence after injection of DOI and DOI+MDL (**Extended Data Fig. E6**). Furthermore, injection of DOI increased 0.5 Hz - 2.0 Hz global coherence. Lastly, across all compound conditions, hemoglobin and calcium exhibited a lag-lead relation, reflected in the positive slope in the phase plots at frequencies below ~1 Hz (**Extended Data Fig. c**).

*N1.2) Regional cross-covariance analysis*

Examination of regional CCF and MSC were computed to assess the topography of compound-specific changes in NVC (**Extended Data Fig. 8**). Before injection of any compound, covariance was strongest in somatosensory and parietal regions and weakest in frontal, cingulate, retrosplenial, and motor regions (**Extended Data Fig. 8a**, left). As evident in post-injection DOI regional HRFs (**Extended Data Fig. 8a**), DOI decreased positive-lag coupling strength across the cortex. Moreover, as observed in the DOI global CCFs (**Extended Data Fig. 8a**), all DOI regional CCFs exhibited a sharp notch at zero lag. Notably, the deepest notches were in secondary motor, retrosplenial, and visual regions. Moreover, DOI CCFs decayed faster in all cortical regions, with stronger decreases in the cingulate and retrosplenial areas. DOI+MDL predominantly reflected pre-injection CCFs, except for negative-lagged decreases in somatosensory and parietal regions.

Regional contributions to global MSC were also evaluated (**Extended Data Fig. 8b**). As reported in global measures, before administration of compounds, strong coherence was restricted to low frequencies (<0.5 Hz) and was strongest in somatosensory and parietal regions. Coherence was weakest in frontal, cingulate, motor, and retrosplenial regions. After injection of DOI, the global MSC peak at 0.8 Hz observed in global measures was primarily driven by increased coherence occurring within higher-order brain regions such as cingulate, secondary motor, and retrosplenial cortex. Following injection of DOI+MDL, regional coherence primarily reflected pre-injection characteristics.

Prior to injection, regional phase relations between neuronal and hemodynamic activity were relatively similar over the cortex (**Extended Data Fig. 8c**). However, under DOI, frontal, cingulate, and retrosplenial regions exhibited significantly altered phase relationships and were only partially reversed in the DOI+MDL condition.

*N2) Inter- versus Intra-network functional connectivity alterations*
After injection of DOI, calcium, seed-based ISA RSFC (**Fig. 6**, left columns) exhibits increased intra-network connectivity in frontal and cingulate (anterior cingulate within the human DMN) regions, but exhibits decreased intra-network connectivity within the retrosplenial (posterior cingulate within the human DMN), visual and M2 networks. Inter-network connectivity increased between frontal and cingulate and between M2 and anterior somatomotor and medial retrosplenial but decreased between cingulate, somatomotor and visual networks. Interestingly, RSFC decreased within the anterior (cingulate) and posterior (retrosplenial) parts of the mouse DMN. After the injection of DOI, silhouette scores (**Extended Data Fig. 10e**, top) consistently decreased across the cortex, indicating a loss of network integration. The largest decreases were in somatomotor and retrosplenial regions. Importantly, MDL administered with DOI did not fully reverse many changes in RSFC from DOI alone (e.g., somatomotor decreases). However, DOI+MDL was associated with increased scores in the lateral visual, posterior somatomotor, and cingulate regions.

Hemodynamic estimates of RSFC (**Fig. 6**, center columns) did not reveal increased intra-network connectivity but did reveal decreased inter-network connectivity between frontal and retrosplenial regions, a phenomenon opposite of the calcium-based RSFC estimates. Silhouette scores decreased in posterior cingulate and increased in medial parts of visual cortex (**Extended Data Fig. 10e**, middle). These changes, contrasted with the changes in the saline group exhibit increased silhouette scores across the cortex (excluding posterior somatomotor and medial retrosplenial regions). MDL did not fully reverse DOI-specific features.

Calcium delta-band RSFC reveals a richer array of changes that are inaccessible to hemodynamic measures of neurophysiology. Specifically, consistent with post-injection DOI ISA calcium RSFC, retrosplenial-cingulate connectivity is decreased; intra-network connectivity is increased in the cingulate region but decreased within the retrosplenial region. In contrast, the decrease in inter-network connectivity between the retrosplenial cortex and the rest of the cortex is more extensive in spatial extent compared to RSFC evaluated over ISA. Silhouette scores increased in the frontal and retrosplenial cortex, and moderately changes around SM borders, indicating a small shift in border locations, as opposed to gross alterations in intra- versus inter-network connectivity.

### ***Supplemental Discussion***

*D1) Post-stimulus undershoots*
DOI was observed to alter the post-stimulus undershoot of task-evoked hemodynamic activity and HRFs. This aspect of the HRF has been attributed to delayed vascular compliance, sustained increases in $CMRO_2$, and/or post-stimulus reductions in cerebral blood flow [13-15]. More recent work has reported that post-stimulus undershoots in hemodynamic responses are due to (and modulated by) post-stimulus neuronal activity [16, 17]. For example, increased inhibitory activity can reduce cerebral blood flow [18-20]. Psychedelic 5-$HT_{2A}$R agonism can result in dose-dependent vasoactive changes [21, 22], which may alter recovery times of vascular compartments. Because MDL alone or in combination with DOI did not alter the post-stimulus undershoot, we speculate that the post-stimulus undershoot observed following DOI depend on receptor activity and signaling pathways agonized by DOI and antagonized by MDL (*e.g.*, 5-$HT2_{A/B/C}$).

*D2) Variance explained differs between task and resting-state conditions*
In line with prior work, the variance explained by Thy1-based activity on local hemodynamics meets or exceeds prior reports [23-25]. Using an updated model of a causal, linear-shift invariant system (*i.e.*, weighted least squares deconvolution), prior to any compound injection, our model fit explains >65%

and >81% variance under resting-state and task conditions, respectively (**Fig. S5**). These differences can likely be linked to changes in the brain's energy demands for processing sensory input compared to those required for sustaining ongoing spontaneous activity [26, 27]. Since our models were fit to each individual stimulus presentation, rather than after block-averaging, task-based activity can be interpreted as superimposed onto resting activity, increasing the signal-to-noise ratio during the periods of stimulus delivery [28], which, in turn, led to an increase in the variance explained.

Figures and Tables:

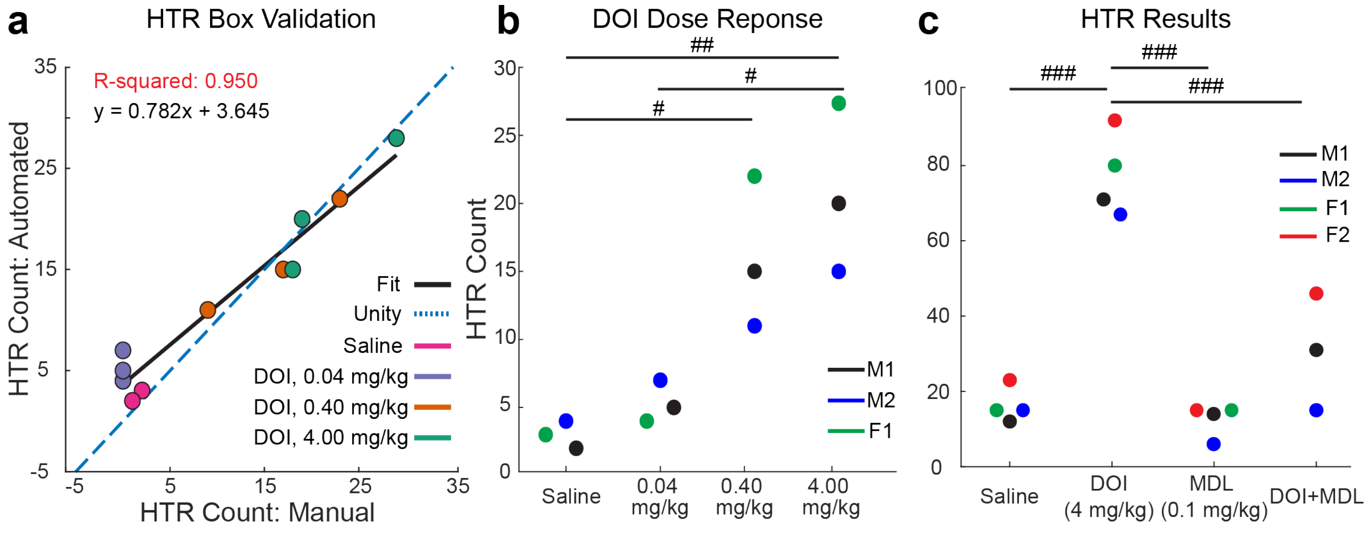


**Supplemental Figure 1. a) Automated assessment of head twitch responses (HTRs).** A custom-built magnetometer was used for all HTR recordings. A magnet placed on the mouse ear induces an electromotive force in the solenoid during an HTR, which is recorded following amplification. **a) Validation of automated HTR analysis.** Automated HTR recordings were validated through comparison to manual scoring of HTRs; validation was performed following saline injection and over a range of log-spaced doses of DOI. Correspondences between manual- and automated- scoring of HTRs were excellent ($R^2 = 0.950$). HTRs were recorded for 10 minutes. **b**) **Dose response of DOI.** HTRs between different doses of DOI were significantly different (ANOVA, p=0.0029; t-test post-hoc with Bonferroni correction, saline vs. 0.40mg/kg, p=0.0406; saline versus 4.00 mg/kg, p=0.0062; 0.04 mg/kg vs. 4.00 mg/kg, p=0.0144). A DOI dose of 0.04 mg/kg did not significantly differ from saline (saline versus 0.04 mg/kg, p>0.99) and was selected for sub-hallucinogenic (low-dose) DOI experiments. **c**) **HTR of compounds.** Saline, DOI, DOI+MDL, and MDL differentially elicited HTRs, with DOI being the only compound to elicit an HTR (ANOVA, p<0.0001; post-hoc t-test with Bonferroni correction, saline vs. DOI, p<0.001; DOI vs. MDL, p<0.001; DOI vs. DOI+MDL, p=0.0002). HTRs were recorded for 30 minutes. #: p<0.05, ##: p<0.01, and ###:  p<0.001.

**Supplemental Table 1.** Optical properties used for WFOI analysis.

| <u>Optical Properties</u> | ***E* ($cm^{-1}$ $M^{-1}$)** | **$\mu_a$ ($cm^{-1}$)** | **$\mu_s'$ ($cm^{-1}$)** | **$X$ (cm)** |
|---|---|---|---|---|
| **Green Excitation LED w/o filter** | $3.4\times10^4$<br>$3.7\times10^4$ | 6.1 | 36 | 0.072 |
| **jRGECO1a Emission (peak)** | $1.6\times10^4$<br>$2.1\times10^4$ | 3.1 | 32 | 0.099 |
| **625nm w/ filter** | $5.5\times10^2$<br>$4.7\times10^3$ | 0.30 | 29 | 0.30 |
| **525nm w/ filter** | $3.5\times10^4$<br>$3.8\times10^4$ | 6.2 | 37 | 0.070 |

Extinction coefficient (*E*), absorption coefficient ($\mu_a$), reduced scattering coefficient ($\mu_s'$) and differential path length ($X$) are tabulated for excitation/emission spectra of jRGECO1a and diffuse reflectance of each LED. Optical properties reported represent a spectrum-weighted average. Top and bottom values for the extinction coefficient represent those for HbO and HbR, respectively.

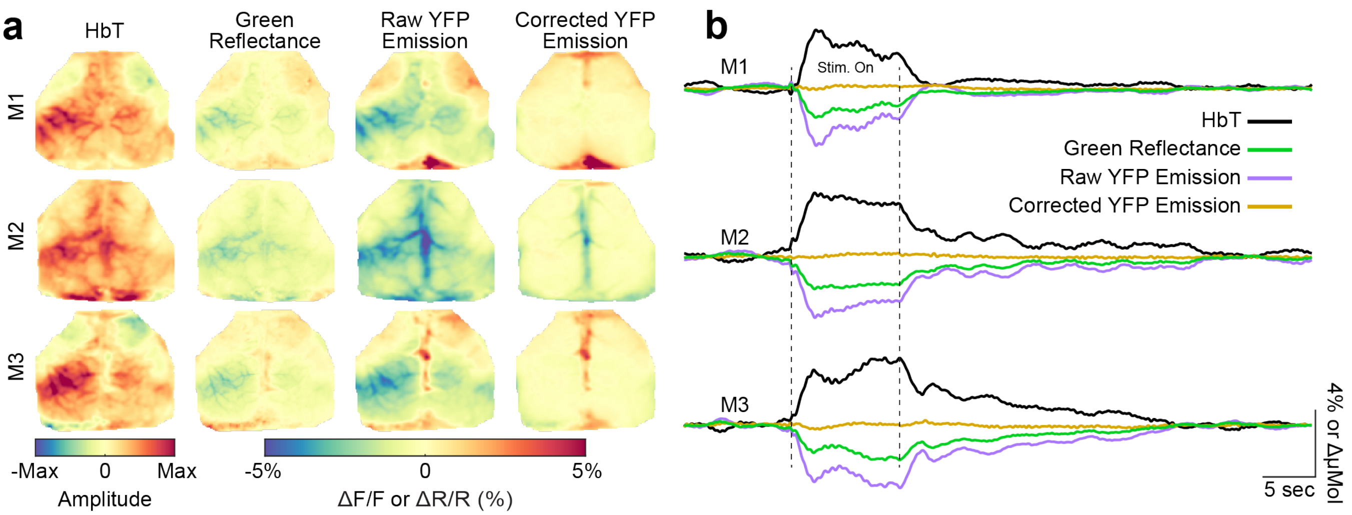


**Supplemental Figure 2. Validation of hemodynamic correction in wide-field calcium fluorescence measurements.** Three Thy1-YFP mice (2M, 1F) were imaged to evaluate the efficacy of our hemodynamic correction algorithm. Mice were subjected to the same procedures and whisker-stimulation paradigm as the experimental mice. **a**) **Topography of hemodynamic correction algorithm.** Maps of HbT, green reflectance, raw YFP-emission, and corrected YFP emission during responses 5-10 seconds) were computed for each mouse. Before hemodynamic correction, response maps of raw YFP emission demonstrate apparent activity during stimulation (decreased $\Delta F/F$), a phenomenon resulting from increased absorption owing to increased blood volume (as reported by changes in green reflectance). After correction, hemodynamic crosstalk is effectively eliminated over most of the cortex. **b**) **Temporal dynamics of hemodynamic correction algorithm.** Time traces within primary somatosensory barrel cortex were averaged together for each mouse. Following correction, covariance between green reflectance and YFP signals significantly decreased after correction (from 0.547$\pm$0.167 to 0.019$\pm$0.012, paired t-test, p=0.0057). Additionally, the sum of the absolute changes in reflectance (*i.e.*, $\frac{\Delta \mathrm{F}}{\langle \mathrm{F} \rangle}$) over the stimulus period significantly decreased after correction (from -268.6$\pm$11.1% to 4.6$\pm$5.0%, paired t-test, p<0.001).

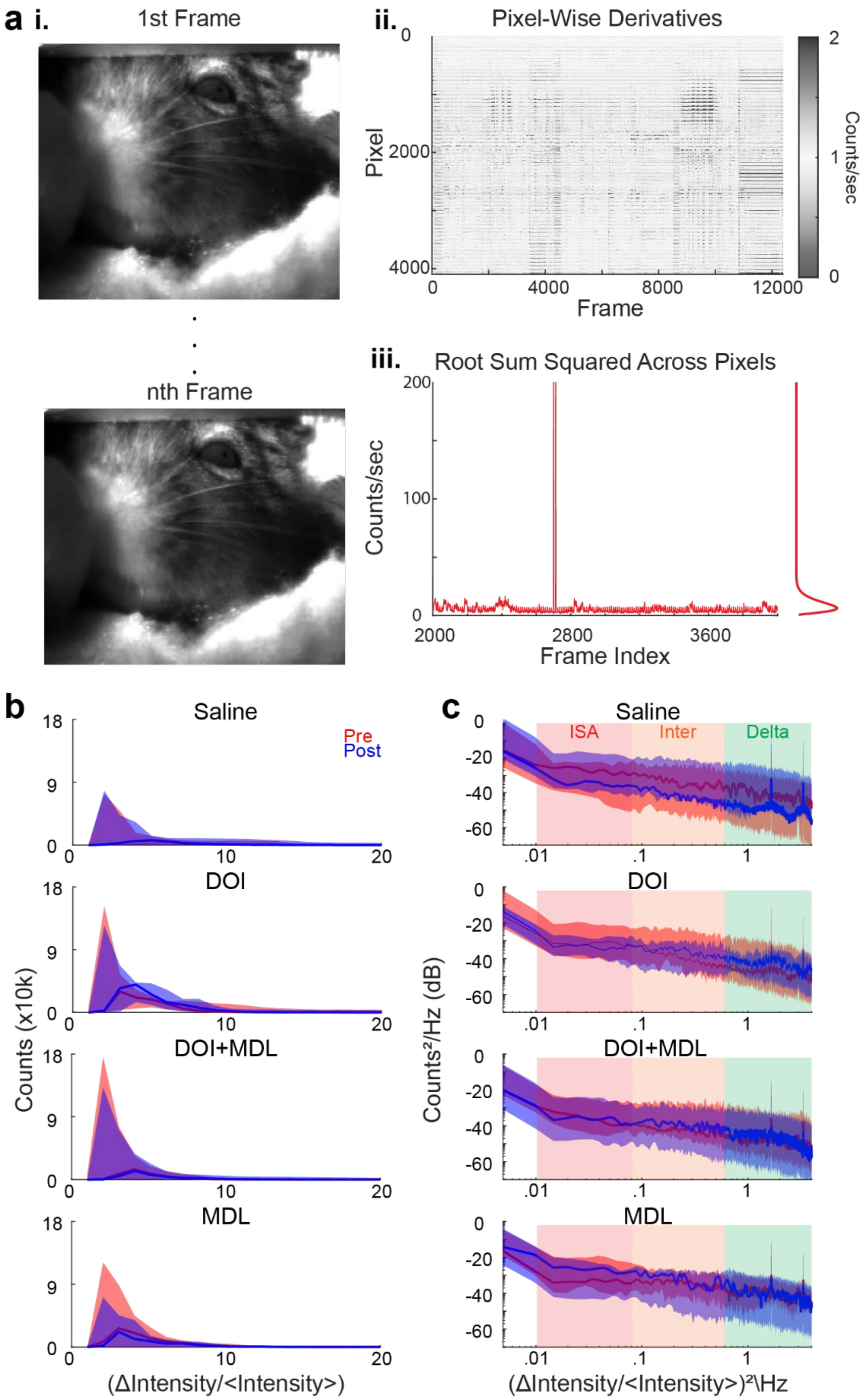


**Supplemental Figure 3. Alternative method to assess movement during WFOI also reveals no differences in movement between compounds.** A CMOS camera (frame rate: 20 Hz, field-of-view ~7cm-by-7cm), was placed in of the mouse to monitor facial and body movement. Frontal illumination was delivered through a 750 nm, near infrared (NIR), LED, and NIR polarizers were used to minimize specular reflection. Movies were recorded and time locked to WFOI. **a) Analysis pipeline of movement quantification**. Movies (**i**) were binned by a factor of two to allow for faster processing time and each pixel was normalized to its mean average intensity over the duration of acquisition. Each pixel was differentiated across time (**ii**), rectified through squaring, summed over pixels, and then the square root was calculated to get a single root-sum squared (RSS) trace of movement for each scan (**iii**). Histograms of RSS traces were averaged across all scans for each mouse and compound group. The spectral content of RSS traces was calculated using a smoothing estimator (pwelch). **b) Histograms of movement traces.** No significant differences were found between pre- and post- distributions, and differences in distributions between pre- and post- injection (post- minus pre-) were not significantly different across compounds. **c) Power spectral density estimates of movement traces.** The spectral content of RSS traces was examined using a smoothing estimator and band-limited integrated power was compared over the infraslow (ISA, 0.01-0.08 Hz), intermediate (0.08-0.50 Hz) and delta-bands (0.5-4.0 Hz). Across frequency bands, no significant differences were found between pre- and post- distributions or across compound. The number of censored runs per 30-minute session were not significantly different between compounds.

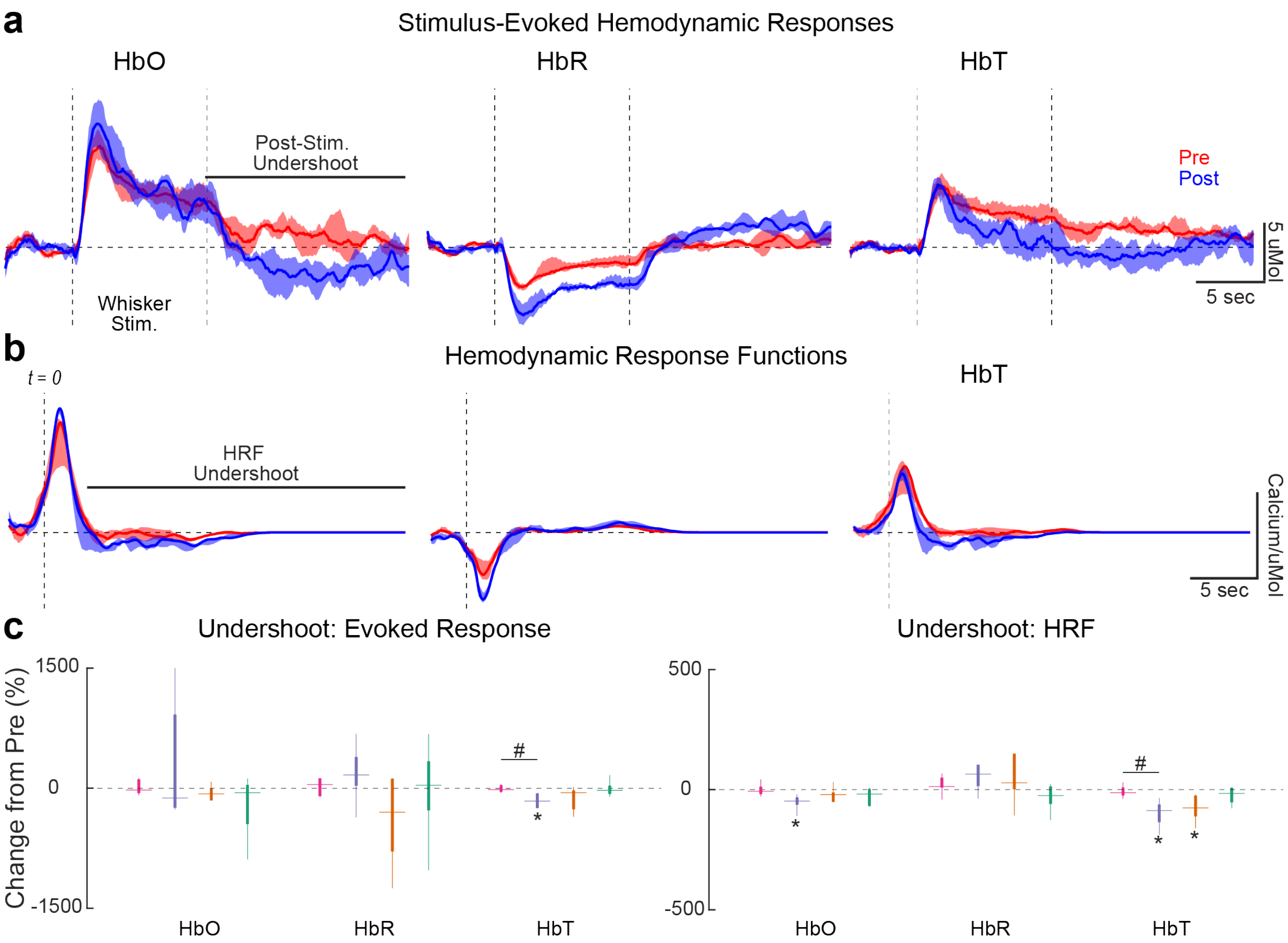


**Supplemental Figure 4) Post-stimulus undershoot is increased under DOI. a) Measured stimulus responses of HbO, HbR, and HbT during whisker stimulation before and after injecting DOI.** Post-stimulus undershoots are larger in all hemoglobin species and are quantified in panel c. **b) HRF estimates using weighted deconvolution for all hemoglobin species before and after DOI.** Data are presented as medians and $25^{th}$ and $75^{th}$ percentiles across mice**. c) Quantification of response undershoot**. The hemodynamic response undershoot (the sum after stimulus-offset; indicated by black dotted line) and HRF undershoot (sum after $t = 2$ sec; indicated by black dotted line) were computed. DOI decreases the hemodynamic response undershoot compared to saline (Kruskal-Wallis: p=0.042; saline versus DOI: p=0.041). Quantification of undershoot changes in oxy-, deoxy-, and total-hemoglobin reveals that HbT undershoot results from decreased oxygenation (-121% [-243% 9%]) and increased deoxygenation (166% [-390%, -30%]). Similar observations occurred in estimated HRFs for each hemoglobin species (Kruskal-Wallis: p=0.003; saline versus DOI: p=0.021; DOI versus MDL: p=0.021). All outliers (>$99.3^{rd}$ percentile) were excluded for visualization purposes only. Differences (pre- versus post-injection) were assessed through Wilcoxon signed-rank tests. *<0.05, **<0.01, ***<0.001. Differential effects of compound were tested using a one-way Kruskal-Wallis and post-Hoc Wilcoxon's sign-rank tests. #<0.05, ##<0.01, ###<0.001. Multiple comparisons were corrected for using the Bonferroni method.

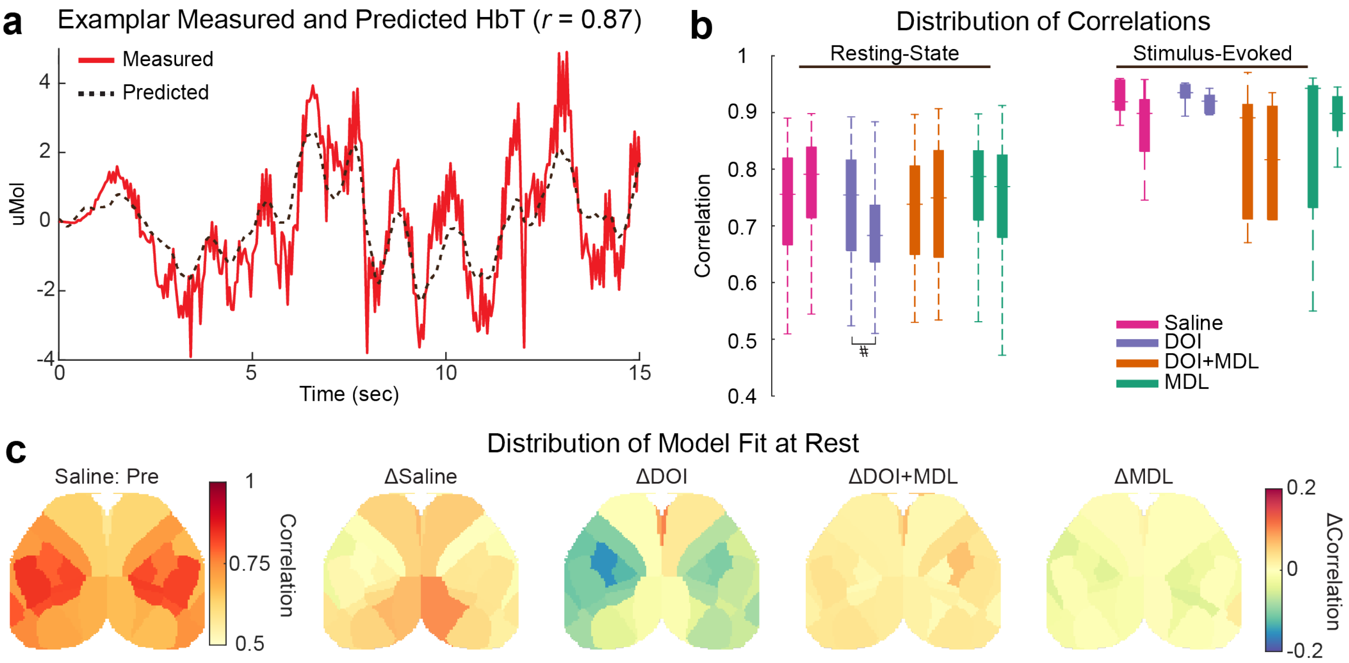


**Supplemental Figure 5). Model fit of neurovascular coupling**. Neurovascular coupling was estimated using weighted, least squares deconvolution between measured calcium (input) and hemodynamic (output) activity. Model fit was evaluated as the Pearson correlation strength between measured and predicted fluctuations in total hemoglobin for each region. **a**) **Predicted HbT corresponds with Measured HbT.** Example time traces of measured (red) and predicted (dotted black) resting-state hemodynamic activity under within the parietal region; the excellent model fit ($r = 0.87$) indicates strong correspondence between measured and predicted signals. **b**) **Distribution of model fit.** The distribution of cortical-averaged correlations between predicted and measured hemodynamics for each group and compound under both resting-state and stimulus-evoked paradigms. Correlations frequently exceed 0.65 across all observations, indicating that hemodynamic activity complies with a linear, causal, and shift-invariant model. *p<0.05. **c**) Cortical topographies of model fit. ***Left***: Before saline injection, correlation strengths are consistently above 0.5. ***Right***: Saline, DOI, DOI+MDL, and MDL minimally altered model fits, however, DOI injection slightly decreased the correlation across the cortex. **c**) **Regional variations in model fit.** Before the injection of any compound, model fit exhibited regional variation, with the largest model fit in somatomotor regions and the lowest in cingulate and secondary motor regions. After the injection of compounds, the model fit increased for all compounds and regions, except DOI. DOI decreased the model fit in the somatomotor and medial/posterior regions and increased the model fit in cingulate cortex.
